# SLFN11-mediated tRNA regulation induces cell death by disrupting proteostasis in response to DNA-damaging agents

**DOI:** 10.1101/2025.01.08.632070

**Authors:** Yuki Iimori, Teppei Morita, Takeshi Masuda, Shojiro Kitajima, Nobuaki Kono, Shun Kageyama, Josephine Galipon, Atsuo T. Sasaki, Akio Kanai

## Abstract

DNA-damaging agents (DDAs) have long been used in cancer therapy. However, the precise mechanisms by which DDAs induce cell death are not fully understood and drug resistance remains a major clinical challenge. Schlafen 11 (SLFN11) was identified as the gene most strongly correlated with the sensitivity to DDAs based on mRNA expression levels. SLFN11 sensitizes cancer cells to DDAs by cleaving and downregulating tRNA^Leu^(TAA). Elucidating the detailed mechanism by which SLFN11 induces cell death is expected to provide insights into overcoming drug resistance. Here, we show that, upon administration of DDAs, SLFN11 cleaves tRNA^Leu^(TAA), leading to ER stress and subsequent cell death regulated by inositol-requiring enzyme 1 alpha (IRE1α). These responses were significantly alleviated by SLFN11 knockout or transfection of tRNA^Leu^(TAA). Our proteomic analysis suggests that tRNA^Leu^(TAA) influences proteins essential for maintaining proteostasis, especially those involved in ubiquitin-dependent proteolysis. Additionally, we identified the cleavage sites of tRNA^Leu^(TAA) generated by SLFN11 in cells, and revealed that tRNA fragments contribute to ER stress and cell death. These findings suggest that SLFN11 plays a crucial role in proteostasis by regulating tRNAs, and thus determines cell fate under DDA treatment. Consequently, targeting SLFN11-mediated tRNA regulation could offer a novel approach to improve cancer therapy.

**GRAPHICAL ABSTRACT:** 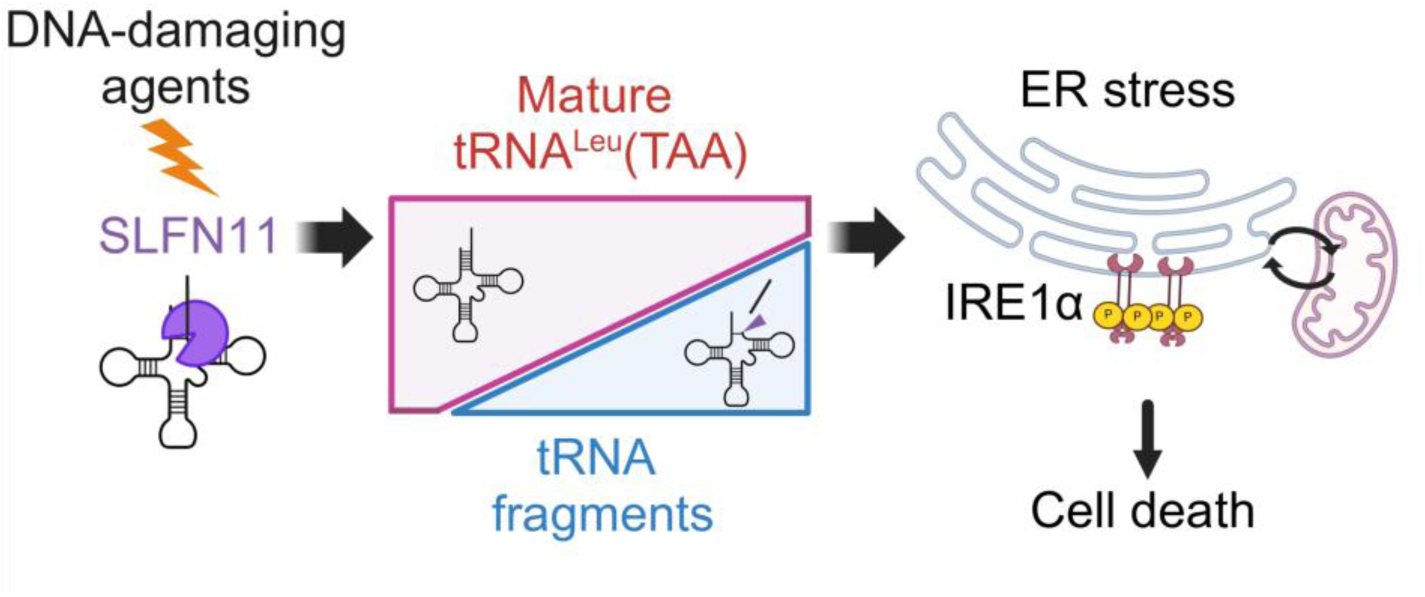

## INTRODUCTION

For decades, DNA-damaging agents (DDAs), such as platinum derivatives, topoisomerase inhibitors, and DNA synthesis inhibitors, have served as first-line treatments for various cancers (1–3). However, the mechanisms by which DDAs induce cell death are not fully understood. Thus, drug resistance and the lack of reliable biomarkers to predict DDA efficacy remain major clinical problems. To address this issue, large cancer cell databases (NCI-60 and Cancer Cell Line Encyclopedia) have been analyzed to identify genes correlated with the sensitivity to DDAs in terms of mRNA expression levels, and *SLFN11* was identified as the gene with the strongest correlation (4,5). SLFN11 was subsequently verified to increase the sensitivity of various cancer cell lines to DDAs, and it is emerging as an important factor with the potential to overcome drug resistance (6). Elucidating the mechanism by which SLFN11 induces cell death is expected to advance new treatment strategies for various cancers.

*SLFN11* is an interferon-inducible gene that inhibits viral replication (7–9). In this process, SLFN11 regulates tRNA levels to inhibit the translation of viral proteins. Similarly, SLFN11 regulates tRNAs in response to DDAs. Upon DDA treatment, SLFN11 cleaves specific tRNAs (particularly type II tRNAs with a V-arm structure, such as tRNA^Leu^ and tRNA^Ser^) and binds to single-stranded DNA, inducing cell death and blocking DNA replication (10–15). Likewise, SLFN12 also cleaves tRNA^Leu^(TAA) and induces cell death (16). The mechanism by which tRNA cleavage induces cell death has been reported to involve the downregulation of tRNA^Leu^(TAA), leading to ribosome stalling at UUA codons and subsequent ribotoxic stress signals (17) or a reduction in proteins with high UUA codon usage frequency, such as ataxia telangiectasia and Rad3-related (ATR) (11). However, the previous studies on UUA codon usage frequency have primarily focused on the proteins involved in DNA damage response and repair mechanisms (11), leaving the broader role of tRNA^Leu^(TAA) in DNA damage and its role in inducing cell death incompletely understood. Furthermore, the effect of tRNA fragments (tRFs) produced by SLFN11 on cell death has not been evaluated.

The level of tRNA is adjusted to match the codon usage of mRNA, and an imbalance can lead to changes in protein expression levels or the production of misfolded proteins (18). The accumulation of misfolded proteins induces endoplasmic reticulum (ER) stress. In most cases, ER stress is alleviated by the unfolded protein response (UPR), which maintains protein homeostasis (proteostasis) by accelerating protein degradation and repressing translation (19). In addition, the ER and mitochondria coordinate to maintain proteostasis through calcium ion transfer and ATP production (20). However, if ER stress persists, the UPR triggers cell death through specific signaling molecules, such as C/EBP homologous protein (CHOP) and the Inositol-requiring enzyme type 1α (IRE1α)-c-Jun N-terminal kinase (JNK) pathway (21,22). Furthermore, tRNA fragments (tRFs) produced by the cleavage of mature tRNAs regulate protein translation and cell death (23–26).

In this study, we investigated how the downregulation of tRNA^Leu^(TAA) leads to cell death under DDA treatment by examining the impact of tRNA^Leu^(TAA) transfection on global protein expression levels. We further explored the role of tRNA^Leu^(TAA) under DDA treatment by analyzing proteins that frequently use UUA codons. Our findings revealed that a decrease in tRNA^Leu^(TAA) induces ER stress and subsequent cell death regulated by IRE1α, and that proteins with high UUA codon usage frequency are often important for proteostasis. Additionally, for the first time, we evaluated the effect of the tRFs produced by SLFN11 on cell death, finding that these tRFs contribute to ER stress and cell death under DDA treatment. Our findings reveal a novel role for SLFN11 in linking DDA treatment to ER stress-related cell death through tRNA regulation.

## MATERIAL AND METHODS

### Cell culture

Ovarian endometrioid adenocarcinoma (TOV-112D) cells (RRID: CVCL_3612) were purchased from the American Type Culture Collection (Manassas, VA, USA). The SLFN11-KO TOV-112D cells were generated in another study (27). Cells were cultured in Dulbecco’s modified Eagle medium (Nacalai Tesque, Kyoto, Japan) supplemented with 10% fetal bovine serum (Biowest, Nuaille, France) and 1% penicillin-streptomycin-amphotericin B (Nacalai Tesque) at 37 °C in 5% CO_2_.

### Cell viability assay and apoptosis assay

TOV-112D cells (2.5 × 10^3^) were seeded onto 96-well white plates (Bio Medical Science, Tokyo, Japan) in 100 µL of medium per well. After incubation for ≥24 hours, the cells were continuously exposed to camptothecin (CPT), a topoisomerase I inhibitor, and cell viability and apoptosis assays were performed.

For the cell viability assay, intracellular ATP levels were measured using the ATPlite 1-step kit (PerkinElmer, Connecticut, USA). After 24 hours of CPT treatment, ATPlite solution (30 µL) was added to each well of a 96-well plate, the cells were lysed, and the luminescence signal from the reaction of intracellular ATP with luciferase in the regent was measured using a plate reader (Infinite M200, TECAN, Männedorf, Switzerland; or Synergy LX, BioTek, Winooski, VT, USA). The ATP level in untreated cells was defined as 100%. Viability (%) of treated cells was defined as ATP in treated cells/ATP in untreated cells × 100.

In the apoptosis assay, phosphatidylserine, exposed on the cell membrane surface, a marker of early apoptosis, was detected using the RealTime Glo Annexin V Apoptosis Assay Kit (Promega, Madison, WI, USA). Briefly, the medium was replaced with 50 µL of medium containing CPT and 1× Detection Reagent, and the plate was incubated at 37 °C in 5% CO_2_. The luminescence signal generated by the binding of Annexin V-luciferase fusion protein to phosphatidylserine was continuously measured without cytolysis. Measurements were performed every 2 hours for up to 24 hours using a plate reader (Infinite M200, TECAN). Values are relative to the mean value of the control samples treated with 0.1% dimethyl sulfoxide (DMSO).

### Northern blotting

Total RNAs were isolated using the Monarch Total RNA Miniprep Kit (New England Biolabs, Ipswich, MA, USA) according to the manufacturer’s protocol. Purified RNAs were denatured by heating at 93 °C for 2 min in formamide gel-loading buffer. The RNA samples (5 µg/lane) were resolved by 10% polyacrylamide gel electrophoresis in the presence of 7 M urea, and transferred onto a nylon membrane (Roche, Basel, Switzerland) for 45 min in 0.5× Tris/borate/ethylenediaminetetraacetic acid (TBE) buffer at 100 mA using a Mini Trans-Blot Cell (Bio-Rad Laboratories, Hercules, CA, USA). After transfer, the RNA samples were fixed to the membrane using FUNA-UV-LINKER (Funakoshi, Tokyo, Japan), and visualized using a digoxigenin (DIG; Roche) detection system according to the manufacturer’s protocols. The RNA probes were synthesized *in vitro* using the DIG RNA labeling kit (Roche) according to the manufacturer’s protocol. Hybridization was carried out overnight at 50 °C. The hybridized membranes were washed three times at 65 °C for 20 min. The RNAs were visualized using an imaging system (ChimiDoc XRS Plus, Bio-Rad). The sequences of the RNA probes and their corresponding positions on tRNA are listed in Supplementary Table S1.

### Transfection of full-length tRNAs and tRNA fragments

The full-length tRNAs and tRNA fragments were synthesized *in vitro* using the CUGA7 *in vitro* Transcription Kit (Nippon Gene, Tokyo, Japan) according to the protocol. The sequences of the synthesized tRNAs and their corresponding positions on each tRNA are listed in Supplementary Table S2. The synthesized RNAs were resolved by 10% polyacrylamide gel electrophoresis in the presence of 7 M urea, and the gel pieces containing the RNAs of the target size were cut from the gel. The RNAs were eluted from the gel pieces by immersion in elution buffer (20 mM Tris-HCl [pH 7.5], 0.5 M ammonium acetate, 10 mM magnesium acetate, 1 mM EDTA, and 0.1% sodium dodecyl sulfate [SDS]) at 37 °C overnight, and purified by phenol–chloroform extraction and ethanol precipitation. The purified RNAs were dissolved in nuclease-free water, denatured at 95 °C for 2 min, cooled at 22 °C for 3 min, and refolded by incubation at 37 °C for 5 min. Cells were transfected with the synthesized full-length tRNAs and tRNA fragments using Lipofectamine 2000 (Thermo Fisher Scientific, Waltham, MA, USA) for 4 hours according to the manufacturer’s protocol.

### Sequencing of tRNA fragments

To sequence the 5′ and 3′ ends of the tRNA fragments, we used a modified RACE method (28). TOV-112D cells were treated with 100 nM CPT for 10 hours, and total RNAs were isolated as described above. Then, 15 µg of total RNA was resolved by 10% polyacrylamide gel electrophoresis in the presence of 7 M urea and visualized by SYBR Gold staining (Thermo Fisher Scientific). The gel pieces containing tRNA fragments of approximately 70 nucleotides were cut from the gel, and the RNAs were eluted from the gel pieces using the method described above. The eluted RNAs were purified by phenol-chloroform extraction and ethanol precipitation. The purified RNA (100 ng) was treated with RNA 5′ pyrophosphohydrolase (New England Biolabs) to replace the 5′ end of RNA with monophosphate, and then was purified again by phenol‒chloroform extraction and ethanol precipitation. The purified RNA was denatured by heating at 93 °C for 2 min, cooled on ice for 5 min, ligated with T4 RNA ligase 1 (New England Biolabs), and purified by ethanol precipitation. Complementary DNA (cDNA) was synthesized using SuperScript III reverse transcriptase (Thermo Fisher Scientific) using the ligated purified RNAs as templates. After treatment with RNase H (New England Biolabs) to remove the RNA templates, the cDNAs were amplified by PCR using Q5 High-Fidelity DNA Polymerase (New England Biolabs). The PCR products were cloned into plasmid pMiniT 2.0 using In-Fusion Snap Assembly Master Mix (Takara Bio, Kusatsu, Japan). Because the presence of a constitutive promoter sequence in pMiniT 2.0 did not give E. coli colonies, the portion without the constitutive promoter sequence was amplified and used as the cloning vector. Plasmid DNAs were isolated from 66 independent colonies and sequenced. The resulting sequences were mapped to the tDNA sequence of tRNA^Leu^(TAA). The 5′ and 3′ ends were determined from the sequence of the ligation cyclized region. The sequences of the oligonucleotides used for reverse transcription and PCR are listed in Supplementary Table S3.

### Proteomic analysis

Proteins were extracted using phase-transfer surfactant (PTS) buffer containing 12 mM sodium deoxycholate (SDC), 12 mM sodium lauroylsarcosinate (SLS), and 100 mM Tris-HCl (pH 9.0), and then quantified using a bicinchoninic acid (BCA) assay kit. We used 10 µg of protein for digestion. Proteins were reduced and alkylated with 10 mM dithiothreitol and 50 mM chloroacetamide, respectively. Protein solutions were 3-fold diluted with 50 mM ammonium bicarbonate. Proteins were digested with Lys-C for 3 hours followed by trypsin overnight at 37 °C. SDC and SLS were removed from the peptide solution according to the PTS method (29,30). Briefly, an equal volume of ethyl acetate was added to the peptide solution, and the mixture was acidified with trifluoroacetic acid (TFA). The mixture was agitated for 2 min and centrifuged to separate the organic and aqueous phases. The organic phase containing SDC and SLS was removed and dried. The dried peptides were resuspended in 5% acetonitrile containing 0.5% TFA and were purified using StageTip (31). Nanoscale liquid chromatography‒ tandem mass spectrometry was performed using a mass spectrometer (Orbitrap Exploris 480, Thermo Fisher Scientific) equipped with an ultra-high-performance liquid chromatography system (Vanquish Neo, Thermo Fisher Scientific). The flow rate was 300 nL/min, and the injection volume was 5 µL. An analytical column (Nikkyo Technos; inner diameter 75 µm, length 12 cm, packed with 3 µm C18 beads) was used. The data were acquired in the data independent mode. Raw data were converted to the mzML format using MSConvertGUI and the results files were analyzed using DIA-NN 1.8.1 with reference to the UniProt human proteome database.

### Western blotting

Cells were lysed using ice-cold lysis buffer (150 mM NaCl, 50 mM Tris-HCl [pH 8.0], 1% Triton X-100, 2% SDS, 0.5% SDC) containing 1% protease inhibitors and 1% phosphatase inhibitors. The lysates were sonicated, kept on ice for 60 min, centrifuged at 15,000 ×*g* for 15 min at 4 °C, and the supernatant was collected. The samples were mixed with sample buffer solution containing 2-mercaptoethanol (Nacalai Tesque) and boiled at 95 °C for 10 min. The samples were subjected to 8% SDS-polyacrylamide gel electrophoresis and transferred to a polyvinylidene difluoride membrane (Immobilon-P, Merck Millipore, Burlington, MA, USA) using a Mini Trans-Blot Cell (Bio-Rad). The membranes were blocked with blocking buffer for 1 hour at room temperature and immunoblotted with primary antibodies overnight at 4 °C. The membranes were washed with Tris-buffered saline containing Tween (TBS-T), incubated with horseradish peroxidase-conjugated secondary antibodies for 1 hour at room temperature, and washed again with TBS-T. The protein signals were visualized using a mini-imaging system (ImageQuant LAS 4000mini, GE Healthcare Corporation Japan, Tokyo, Japan) with a Chemi-Lumi One L assay kit (Nacalai Tesque). The band intensity was quantified using ImageJ software. The membranes were stripped using a stripping solution to remove the antibody and reblotted with other antibodies. The primary antibodies, secondary antibodies, and blocking buffers are listed in Supplementary Table S4.

### Bioinformatics analysis

Human genome data were downloaded from the NCBI database (accession number: GCF_000001405.40). The codon usage frequencies were calculated using a custom Perl script. The average codon usage frequency was calculated for each gene. For proteomics analysis, data expressed as the total amount of two or more proteins were excluded from the analysis of TTA codon usage frequency. All enrichment analyses were performed using Metascape (32) with the Gene Ontology Biological Processes database. Cluster analysis was performed using Morpheus (https://software.broadinstitute.org/morpheus). Hierarchic clustering was performed using one minus Pearson correlation.

### Statistical analysis and reproducibility

No statistical method was used to predetermine the sample size. Sample sizes were chosen according to common standards. Most experiments were repeated two or three times; the exact number of independent experiments is stated in the figure legend. Data are shown as mean ± standard deviation (SD). One-way or two-way analysis of variance (ANOVA) followed by the appropriate post hoc test for multiple comparisons were performed as appropriate, and the method is described in the figure legends. Significance was accepted at *P* < 0.05. Asterisks indicate the following statistical significance: \**P* < 0.05, \*\**P* < 0.01, and \*\*\**P* < 0.001. The statistical analyses were carried out with GraphPad Prism 10 (GraphPad Software, Boston, MA, USA).

## RESULTS

### SLFN11-dependent downregulation of tRNA^Leu^(TAA) induces cell death in human cancer cells treated with camptothecin

We utilized a system in which cell death was induced in ovarian endometrioid adenocarcinoma (TOV-112D) cells by camptothecin (CPT) (27), a topoisomerase I inhibitor, and examined the induction of apoptosis and alterations in mature tRNA levels or the appearance of tRFs over time. To verify the dependency on SLFN11, experiments were conducted using both parent and *SLFN11*-knockout (KO) cells (Supplementary Figures S1A) (27). In parent cells, CPT induced an apoptotic signature within 24 hours at concentrations above 25 nM (Figure 1A) and significantly reduced cell viability at 24 hours at concentrations above 100 nM (Supplementary Figures S1B). However, the induction of an apoptotic signature or a reduction of cell viability were not observed in *SLFN11*-KO cells at any concentration of CPT (Figure 1B and Supplementary Figures S1B). Based on these results, 100 nM CPT was selected for subsequent experiments, where an apoptotic signature was observed 10 hours after CPT administration in parent cells.

**Figure 1:**
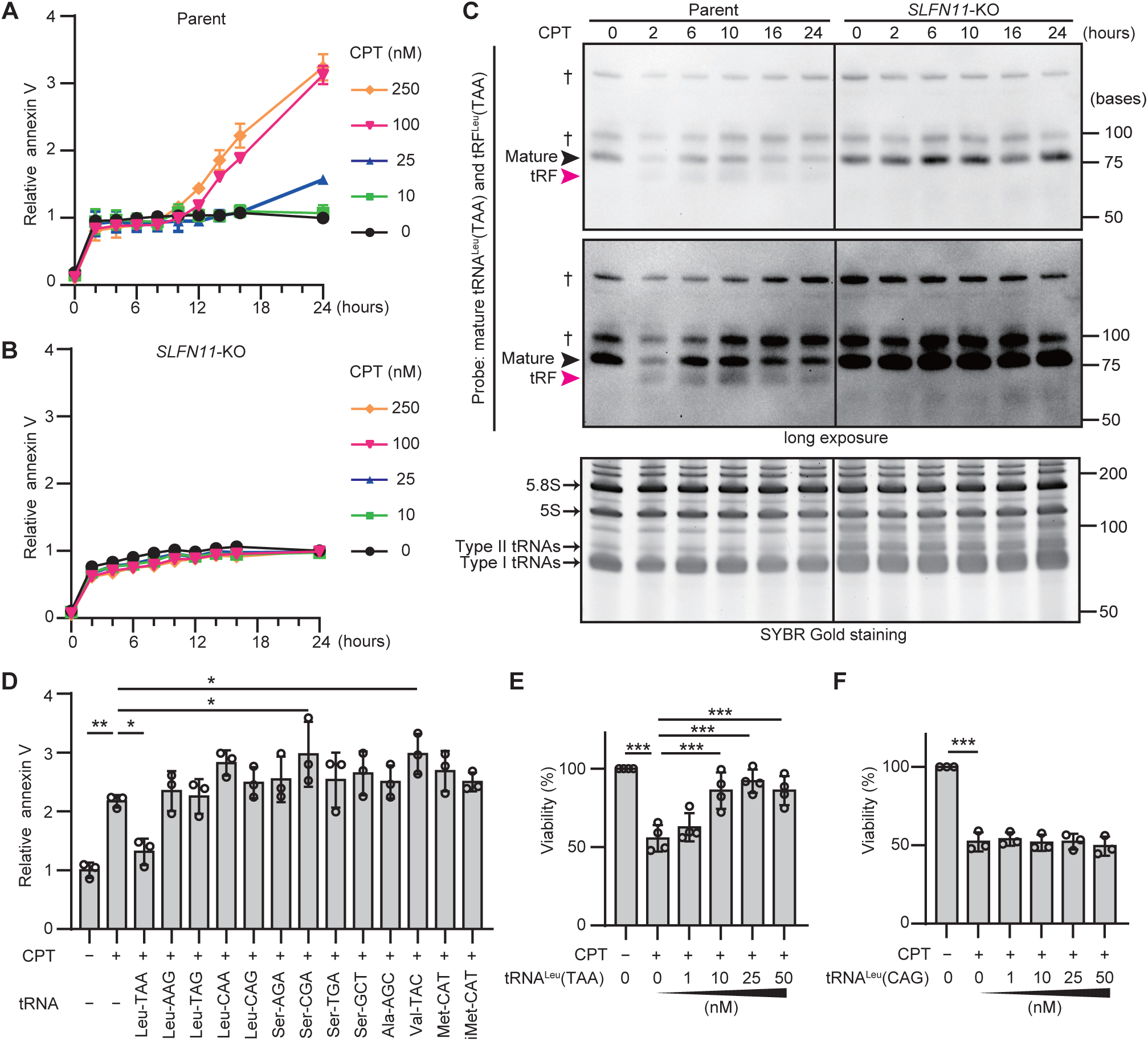
SLFN11-dependent downregulation of tRNA^Leu^(TAA) induces cell death in human cancer cells during CPT treatment. (**A** and **B**) Induction of the apoptotic signature by CPT was quantified at the indicated time points using the annexin V apoptosis assay. TOV-112D parent cells (A) and SLFN11-KO cells (B) were treated with the indicated concentrations of CPT for 24 hours. Values are relative to the mean value of the control samples treated with 0.1% dimethyl sulfoxide (DMSO) for 24 hours. Data are shown as mean ± SD (three technical replicates). The results are representative of two independent experiments. (**C**) Expression levels of mature (black arrowhead) or tRNA fragment (tRF) (red arrowhead) of tRNA^Leu^(TAA) were analyzed by northern blotting. TOV-112D parent cells and *SLFN11*-KO cells were treated with CPT (100 nM) for the indicated times. Daggers indicate the position of the presumed precursor tRNAs. Long exposure data for the upper panel is shown in the middle panel. SYBR Gold staining of the gel shown in the upper panel is presented in the bottom panel. The marker size is indicated on the right side. Representative data from two independent experiments are shown. (**D**) Apoptotic signature of TOV-112D cells assessed after transfection with the indicated full-length tRNA (10 nM) followed by treatment with CPT (100 nM) for 24 hours. (**E** and **F**) Cell viability of TOV-112D cells assessed after transfection with tRNA^Leu^(TAA) (E) or tRNA^Leu^(CAG) (F) at the indicated doses, followed by treatment with CPT (100 nM) for 36 hours. Data are shown as mean ± SD (four biological replicates for panel E, three biological replicates for panel D and F). One-way ANOVA with Dunnett’s multiple comparison tests was used. \**P* < 0.05, \*\**P* < 0.01, and \*\*\**P* < 0.001.

We performed time-series analyses of mature tRNA levels and the appearance of tRFs in cells treated with 100 nM CPT using northern blotting (Figure 1C and Supplementary Figures S2 and S3). These experiments employed 10 probes corresponding to six type II tRNAs [tRNA^Leu^(TAA), tRNA^Leu^(CAG), tRNA^Leu^(CAA), tRNA^Leu^(AAG), tRNA^Ser^(GCT), and tRNA^Ser^(CGA)] and four type I tRNAs [tRNA^Ala^(AGC), tRNA^Val^(TAC), tRNA^Met^(CAT), and tRNA^iMet^(CAT)], which were preliminarily examined in conditions that specifically detected the target tRNAs (Supplementary Figures S2). For probes detecting multiple tRNAs, the total detected amount was evaluated (Supplementary Figures S2D,F,G and S3D,F,G). We detected tRFs of approximately 70 nucleotides in length using all type II tRNA probes in the parent cells (Figure 1C and Supplementary Figures S3A-F). Among type I tRNAs, tRFs were only detected with tRNA^iMet^(CAT) (Supplementary Figures S3G-J). The properties of tRNA^iMet^(CAT), an initiator tRNA, may differ from those of the other type I tRNAs. These tRFs were not detected in the *SLFN11*-KO cells.

Figure 1C presents representative time-series data for tRNA^Leu^(TAA), which showed a suppressive effect on CPT-induced cell death in subsequent experiments. In parent cells, mature tRNA^Leu^(TAA) levels decreased, and tRF^Leu^(TAA) appeared within 2 hours of CPT administration. In *SLFN11*-KO cells, the mature tRNA^Leu^(TAA) levels were higher than those in parental cells after CPT administration, and tRF^Leu^(TAA) was not detected at any time. The levels of 5*S* and 5.8*S* rRNA, used as controls, did not change (Figure 1C, bottom panel). These findings suggest that the downregulation of mature tRNAs and the production of tRFs are early events that may be involved in the initiation of apoptosis.

Previous studies have shown that the downregulation of tRNA^Leu^(TAA) induces apoptosis during CPT treatment (11). In our system, CPT-induced apoptosis was accompanied by mature tRNA downregulation and tRF production in at least six type II tRNAs and tRNA^iMet^. To determine which of the decreased tRNAs was the main factor inducing apoptosis, rescue experiments were performed. We examined whether CPT-induced apoptosis of TOV-112D cells could be suppressed by transfection with full-length tRNAs, using nine type II tRNAs and four type I tRNAs. Only tRNA^Leu^(TAA)-transfected cells exhibited significant suppression of the CPT-induced apoptotic signature (Figure 1D). Furthermore, the CPT-induced reduction in cell viability was suppressed by transfection with tRNA^Leu^(TAA) in a concentration-dependent manner up to 10 nM of tRNA^Leu^(TAA), with the same effect at higher concentrations (Figure 1E). In contrast, transfection of tRNA^Leu^(CAG) did not affect cell viability at any concentration (Figure 1F). These results suggest that downregulation of tRNA^Leu^(TAA) specifically plays a crucial role in inducing apoptosis during CPT treatment, unlike other tRNAs.

### Downregulation of tRNA^Leu^(TAA) induces ER stress and cell death during CPT treatment

Downregulation of tRNA^Leu^(TAA) induces cell death by decreasing the protein expression of ATR (11), which has high UUA codon usage frequency (2.8% of all codons). However, in our experiments, ATR expression did not decrease until 24 hours after CPT administration, while cell death had already occurred (Supplementary Figures S4A). This suggests that other tRNA^Leu^(TAA)-regulated factors may play a more immediate role in CPT-induced cell death.

To identify the factors involved in apoptosis induced by tRNA^Leu^(TAA) downregulation, proteome analysis was performed 10 hours after CPT administration, when apoptotic signatures began to increase. Comparisons were made between the control group without CPT treatment and the CPT-treated group (i.e., the apoptosis-induced state) (Figure 2A and Supplementary Figures S4B), as well as between the CPT-treated group and the CPT+tRNA^Leu^(TAA)-transfected group (i.e., cells rescued from apoptosis) (Figure 2B and Supplementary Figures S4C).

**Figure 2:**
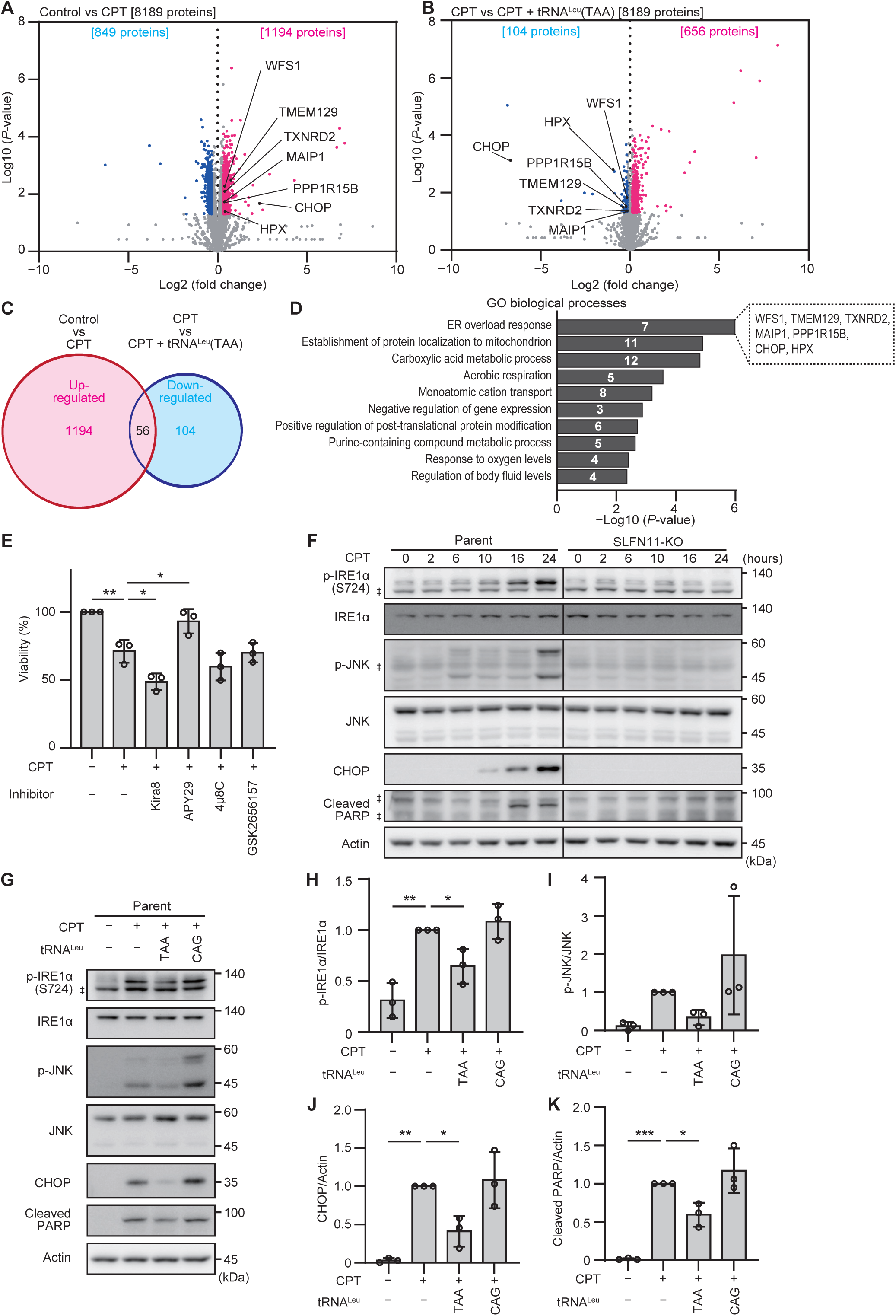
Downregulation of mature tRNA^Leu^(TAA) induces ER stress and causes cell death during CPT treatment. TOV-112D cells were treated with 0.1% DMSO (control), CPT (100 nM), or CPT (100 nM) + full-length tRNA^Leu^(TAA) (10 nM). After 10 hours of treatment, the protein expression levels were evaluated by proteomic analysis (three technical replicates). A total of 8189 proteins were detected. (**A**) Volcano plots of protein expression from TOV-112D cells treated with CPT relative to control cells treated with 0.1% DMSO. A total of 1194 proteins were upregulated (*P* < 0.05 and fold change [FC] > 1.25; red dots) and 849 proteins were downregulated (*P* < 0.05 and FC < 0.8; blue dots). (**B**) Volcano plots of protein expression from TOV-112D cells treated with CPT + full-length tRNA^Leu^(TAA) relative to cells treated with CPT. A total of 656 proteins were upregulated (*P* < 0.05 and FC > 1.1; red dots) and 104 proteins were downregulated (*P* < 0.05 and FC < 0.9; blue dots). The proteins involved in ER overload in (D) are annotated with black letters in (A) and (B). (**C**) Venn diagram of the overlap between upregulated proteins in (A) and downregulated proteins in (B). The total number of overlapping proteins is shown in the center in black letters. (**D**) Enrichment analysis of the overlapping proteins in (C). The white numbers in the bars indicate the number of proteins in each group. The proteins involved in the ER overload response are annotated on the right of the bar. (**E**) Cell viability of TOV-112D cells assessed after 24 hours of treatment with CPT (100 nM) and the stress sensor inhibitors of the UPR, including Kira8 (1µM), APY29 (100 nM), 4µ8c (50 µM), and GSK2656157 (1 µM). (**F**) Western blotting analysis of the time-course of the UPR pathways in TOV-112D parent cells and SLFN11-KO cells after administration of CPT (100 nM). (**G**) Western blotting analysis of the UPR pathway in TOV-112D parent cells transfected with the indicated tRNA (10 nM) followed by treatment with CPT (100nM) for 24 hours. (**H-K**) Quantification of the expression levels of phospho-IRE1α (H), phospho-JNK (I), CHOP (J), and cleaved PARP (K) shown in (G). All data were normalized to the expression levels of IRE1α (H), JNK (I), or actin (J,K) in the same sample, and are relative to the values of samples treated with CPT (100 nM). Representative results from two (F) or three (G) independent experiments are shown. Double daggers indicate the position of the non-specific bands. The marker size is indicated on the right side. In (E and H-K), all data are shown as mean ± SD (three biological replicates). One-way ANOVA with Dunnett’s multiple comparison test was used. \**P* < 0.05, \*\**P* < 0.01, and \*\*\**P* < 0.001.

As a result, 56 proteins were identified that showed increased expression in the apoptosis-induced state and decreased expression in the cells rescued from apoptosis (Figure 2C). The enrichment analysis showed that the proteins involved in the ER overload response (7 proteins) were ranked first, followed by mitochondria-related proteins (Figure 2D). CHOP, a key transcription factor in ER stress-induced apoptosis (33), showed the greatest increase among all ER overload response proteins during CPT treatment (log2 fold change = 2.33). Other proteins involved in maintaining ER and mitochondrial homeostasis were also identified, including those involved in calcium homeostasis (WFS1, MAIP1), ER-associated protein degradation (TMEM129), and regulation of reactive oxygen species (TXNRD2). These results suggest that downregulation of tRNA^Leu^(TAA) triggers ER and mitochondrial stress, leading to apoptosis.

To elucidate the pathway of CPT-induced cell death in TOV-112D cells, we focused on the UPR apoptotic pathway, and assessed the effects of inhibiting the UPR stress sensors using an IRE1α type II kinase inhibitor (Kira8, a mono-selective IRE1α inhibitor that allosterically attenuates IRE1α RNase activity), an IRE1α type I kinase inhibitor (APY29, an allosteric modulator that inhibits IRE1α autophosphorylation while enhancing its RNase activity), an IRE1α RNase inhibitor (4μ8c), and a protein kinase R-like ER kinase (PERK) inhibitor (GSK2656157) (see Supplementary Figure S5A and Supplementary Table S5). CPT-induced cell death was enhanced by Kira8 and partially suppressed by APY29 (Figure 2E and Supplementary Figures S5B-E), indicating that IRE1α plays a regulatory role in CPT-induced cell death. Western blot analysis further verified the activation of the UPR apoptotic pathway under CPT treatment (Figure 2F). In the parent cells, phosphorylation of IRE1α and JNK began 6 hours after CPT administration, after which CHOP was detected at 10 hours, and cleaved poly(ADP-ribose) polymerase (PARP), an apoptotic marker, at 16 hours (Figure 2F). These signals were suppressed by APY29 (Supplementary Figures S5F-J), indicating this pathway is involved in CPT-induced apoptosis. As expected, these UPR signals were absent in *SLFN11*-KO cells at all times (Figure 2F).

To assess the relationship between tRNA^Leu^(TAA) levels and the activation of the UPR apoptotic pathway during CPT treatment, we transfected TOV-112D cells with full-length tRNA^Leu^(TAA) and examined the activation state of the UPR signaling pathway (Figure 2G-K). tRNA^Leu^(CAG) was used as a control to validate the specificity of tRNA^Leu^(TAA). Phosphorylation of IRE1α and JNK, as well as the induction of CHOP and cleaved PARP under CPT treatment, were suppressed by transfection with full-length tRNA^Leu^(TAA), but not by transfection with full-length tRNA^Leu^(CAG). These findings demonstrate that, during CPT treatment, downregulation of tRNA^Leu^(TAA) triggers persistent ER stress, activates the IRE1α‒JNK pathway, induces CHOP expression, and ultimately leads to cell death.

### tRNA^Leu^(TAA) has the potential to regulate proteostasis through translation

To investigate the mechanism by which downregulation of tRNA^Leu^(TAA) induces ER stress, we hypothesized that downregulation of tRNA^Leu^(TAA) impairs the translation of proteins with high UUA codon usage frequency, leading to ER stress. To test this hypothesis, we analyzed the UUA codon usage frequency among proteins positively correlated with tRNA^Leu^(TAA) levels (overlapping proteins in Supplementary Figures S4D) and those inversely correlated (overlapping proteins in Figure 2C) using the proteomics data in Figure 2. Proteins with high UUA codon usage frequency (>2% of all codons) were more prevalent in the positively correlated proteins (16.7%) compared with all detected proteins (10.6%) and were completely absent in the inversely correlated proteins (0%) (Supplementary Figures S6A). These results strongly suggest that the tRNA^Leu^(TAA) levels influence the expression levels of proteins with high UUA codon usage frequency.

To explore the role of proteins with high UUA codon usage frequency, we examined the UUA codon usage frequency among 20,679 proteins listed in the NCBI database. UUA is a rare codon, absent in approximately 25% of human proteins (Figure 3A). Focusing on proteins with UUA codon usage frequency of >3% (313 proteins), enrichment analysis revealed that many of these proteins play critical roles in proteostasis, particularly in processes such as protein transport (e.g., Golgi vesicle transport: 25 proteins) and mitochondrial function (e.g., mitochondrial respiratory chain complex assembly: 26 proteins; mitochondrial RNA metabolic process: 17 proteins) (Figure 3B). Next, using protein data from 148 genes with UUA codon usage frequency of >3% according to the proteomics data in Figure 2, we analyzed which proteins were associated with the changes in tRNA^Leu^(TAA) levels during apoptosis induction and its rescue. Figure 3C shows the results of the clustering analysis of protein level changes in the control group without CPT treatment, the CPT-treated group [apoptosis-induced state with decreased tRNA^Leu^(TAA) levels], and the CPT+tRNA^Leu^(TAA) transfection group (rescued state from apoptosis). As a result, the protein level patterns could be classified into two major clusters.

**Figure 3:**
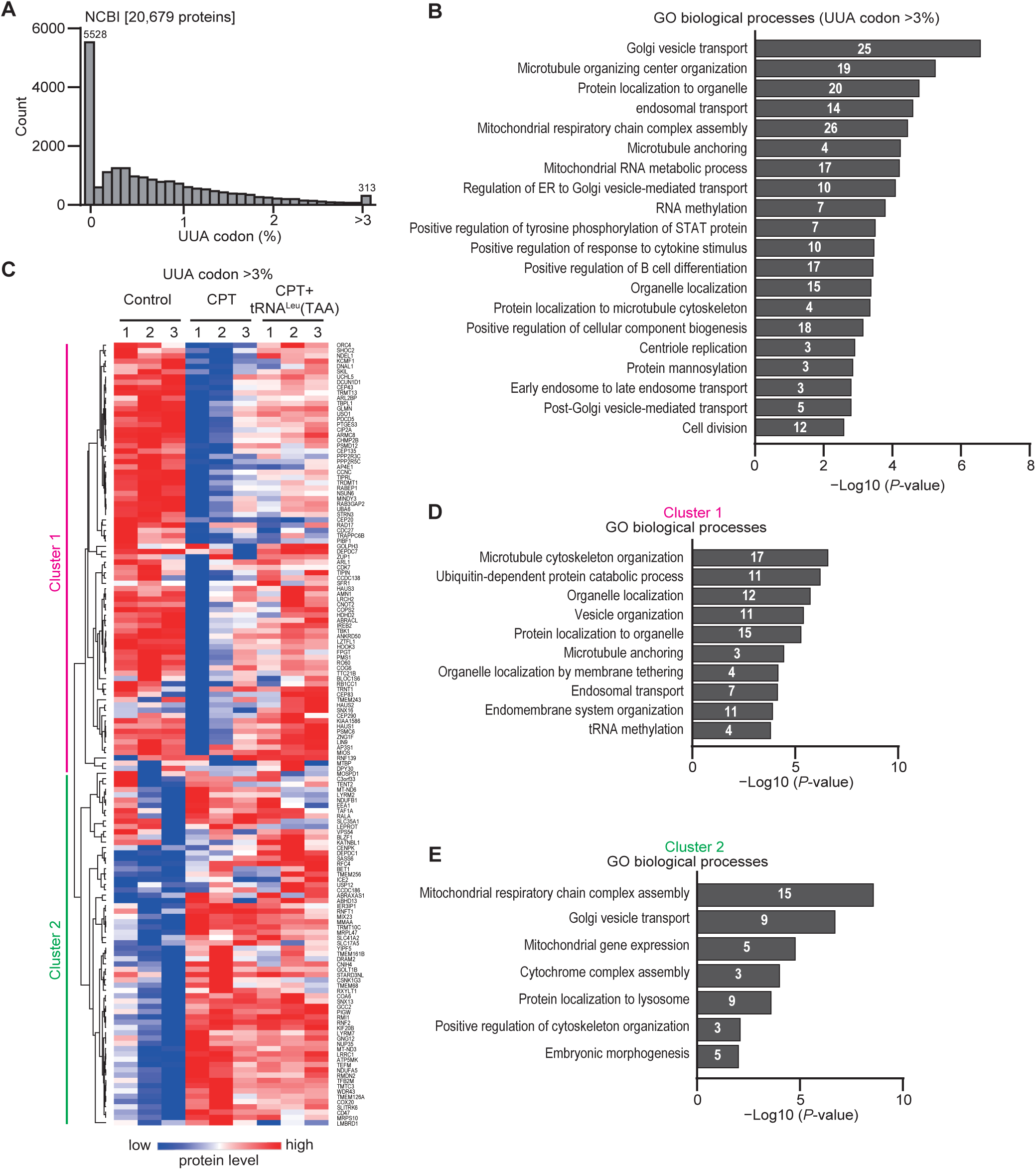
tRNA^Leu^(TAA) has the potential to regulate proteostasis through translation. (**A**) Distribution of UUA codon usage frequency in proteins. A total of 20,679 coding sequences obtained from NCBI were analyzed. (**B**) Enrichment analysis of proteins with a high frequency (>3%) of UUA codon (313 proteins) in dataset (A). (**C**) Heatmap of hierarchic clustering based on one-minus Pearson correlation of the expression level of 148 proteins with a high frequency (> 3%) of TTA codon detected by proteomics analysis. The name of each cluster is shown on the left and the name of each protein is shown on the right. (**D** and **E**) Enrichment analysis of cluster 1 (D) and cluster 2 (E). The white numbers in the bars indicate the number of proteins in each group.

Cluster 1 proteins were correlated with the tRNA^Leu^(TAA) level. Enrichment analysis showed a high ranking of proteins related to microtubule cytoskeleton organization (17 proteins) and ubiquitin-dependent protein catabolic process (11 proteins) (Figure 3D). Notably, the previous analysis also showed that many ubiquitin ligases exhibited significant changes in protein levels correlated with tRNA^Leu^(TAA) levels (Supplementary Figures S4D and E). We investigated ubiquitination after CPT administration and observed a decrease in ubiquitinated proteins in parent cells, which was restored by tRNA^Leu^(TAA) transfection (Supplementary Figures S6B and C). Even when the degradation of ubiquitinated proteins was inhibited using the proteasome inhibitor MG132, CPT treatment reduced the abundance of ubiquitinated proteins in parent cells, suggesting that the production of ubiquitinated proteins is reduced by a reduction in tRNA^Leu^(TAA) levels. One ubiquitin ligase that was significantly reduced after CPT treatment was RCHY1, an essential E3 ubiquitin ligase involved in the degradation of nascent polypeptides during ribosome stalling (34) (UUA codon usage frequency: 2.6%) (Supplementary Figures S4E). These findings suggest that impaired ubiquitin-dependent protein degradation due to reduced tRNA^Leu^(TAA) levels may lead to defective protein accumulation, thereby disrupting proteostasis.

In comparison, Cluster 2 proteins exhibited increased expression during CPT treatment and tRNA^Leu^(TAA) rescue treatment compared to the untreated control samples. Enrichment analysis revealed a high ranking of proteins related to mitochondrial respiratory chain complex assembly (15 proteins) and golgi vesicle transport (9 proteins) (Figure 3E). Although the increase in these proteins suggests a response to CPT, further investigation is needed to clarify whether tRNA^Leu^(TAA) levels influence these processes. This is because reduced tRNA^Leu^(TAA) levels may lead to translation errors or ribosomal frameshifting, potentially resulting in defective protein production (18,35).

Taken together, these results indicate that tRNA^Leu^(TAA) is crucial for the translation of proteins essential to proteostasis, including those involved in protein transport, degradation, and mitochondrial function. This suggests that the downregulation of tRNA^Leu^(TAA) may disrupt proteostasis by causing translation inhibition or errors.

### tRNA fragments generated by SLFN11 promote CPT-induced cell death

The previous experiments focused on the downregulation of mature tRNAs. In this study, we investigated whether SLFN11-dependent tRNA fragments play a role in CPT-induced cell death. We first identified the tRF^Leu^(TAA) cleavage sites in cultured cells using a modified version of the rapid amplification of cDNA ends (RACE) method. Three types of tRF^Leu^(TAA) were identified and named tRF-1, −2, and −3. As shown in the secondary structure of tRF^Leu^(TAA) (Figure 4A), tRF-1 was cleaved at a site 10 nucleotides from the 3′ end, matching prior *in vitro* findings (10). tRF-2 had a four-nucleotide preprocessing sequence at the 5′ end and was cleaved 26 nucleotides from the 3′ end. tRF-3 was cleaved at a site five nucleotides from the 5′ end and 16 nucleotides from the 3′ end. tRF^Lau^(TAA) accounted for 42% of tRF-1, 9% of tRF-2, and 5% of tRF-3. tRF-1 comprised almost half of the sequenced fragments (Supplementary Table S6).

**Figure 4:**
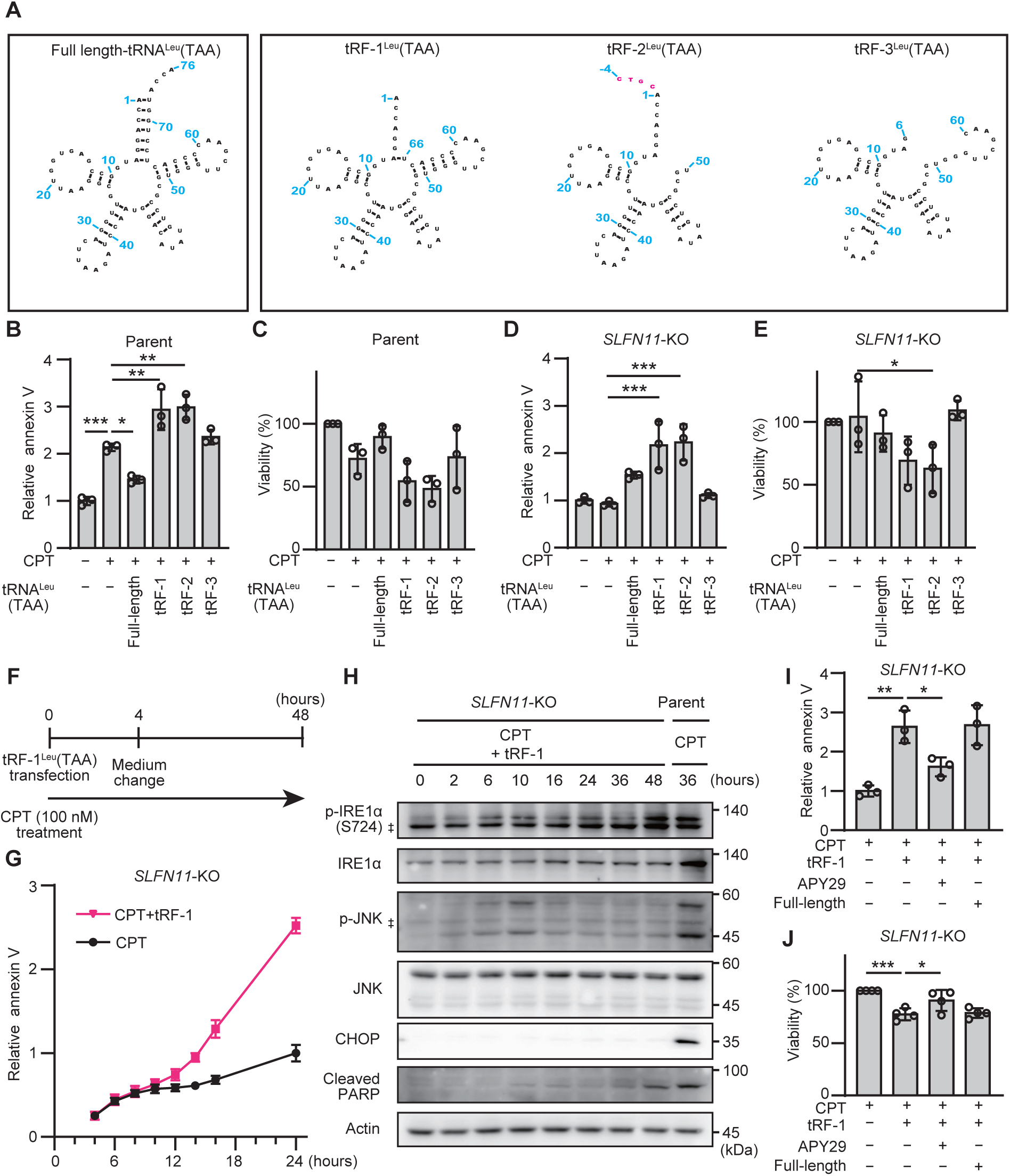
tRNA fragments generated by SLFN11 promote CPT-induced cell death. (**A**) The secondary structure of mature tRNA^Leu^(TAA) and cleavage site of three representative tRF^Leu^(TAA) (tRF-1, tRF-2, and tRF-3) are shown. The positions of the nucleotides are indicated by blue numbers according to the universal tRNA positioning rules (36). Pink letters indicate the 5′ leader sequences before processing. (**B-E**) Induction of cell death by tRNA fragments. The apoptotic signature (B and D) and cell viability (C and E) of TOV-112D parent cells (B and C) or SLFN11-KO cells (D and E) were assessed after 24 hours of treatment with CPT (100 nM) following transfection with full-length tRNA^Leu^(TAA) or tRF^Leu^(TAA)-1, −2, or −3 (10 nM). (**F**) Schematic of the time-course analysis of TOV-112D SLFN11-KO cells treated with tRF-1 (25 nM) and CPT (100 nM). (**G**) Apoptotic signature quantified in real time up to 24 hours using the Annexin V apoptosis assay. Values are relative to the mean value of the control samples treated with CPT (100 nM) for 24 hours. Data are shown as the mean ± SD (three technical replicates). The results are representative of two independent experiments. (**H**) Western blotting analysis of the time-series of the UPR pathway. Representative results from two independent experiments are shown. As a positive control, samples treated with CPT (100 nM) for 36 hours in TOV-112D parent cells are placed in the far-right column. Double daggers indicate the position of the non-specific bands. The marker size is indicated on the right side. (**I**) Apoptotic signature of TOV-112D SLFN11-KO cells assessed 24 hours after treatment with APY29 (100 nM) or full-length tRNA^Leu^(TAA) (10 nM) in addition to CPT (100 nM) and tRF-1^Leu^(TAA) (25 nM). (**J**) Cell viability of TOV-112D SLFN11-KO cells assessed 48 hours after treatment using the same conditions shown in (I). Values are relative to the values of control samples treated with 100 nM CPT. All data in (B-E,I,J) are shown as the mean ± SD (three biological replicates for B-E,I; four biological replicates for J). One-way ANOVA with Dunnett’s multiple comparison test was used. \**P* < 0.05, \*\**P* < 0.01, and \*\*\**P* < 0.001.

To analyze the ability of tRF-1, −2 and −3 to induce cell death, cells were transfected with tRF-1, −2, or −3 during CPT treatment. tRF-1 and tRF-2, but not tRF-3, enhanced or induced cell death during CPT treatment in both parent cells (Figure 4B and C) and *SLFN11*-KO cells (Figure 4D and E). These results suggest that specifically shaped tRNA fragments can induce cell death independently of SLFN11.

Next, we evaluated the potential role of tRF-1 in CPT-induced cell death by treating *SLFN11*-KO cells with CPT from the same time as tRF-1 transfection (Figure 4F) and analyzing the apoptotic signature and UPR signaling. The apoptotic signature started to increase 10 hours after treatment (Figure 4G). Furthermore, IRE1α and JNK were gradually phosphorylated from 2 to 10 hours (Figure 4H). Cell death induced by tRF-1 was partially suppressed by APY29, an IRE1α type I kinase inhibitor (Figure 4I and J). These responses were similar to those observed in the parental cells treated with CPT. However, CHOP was not detected (Figure 4H), and full-length tRNA^Leu^(TAA) did not suppress tRF-1-induced cell death (Figure 4I and J). These results suggest that tRF-1 could play a role in CPT-induced cell death by activating IRE1α, without antagonism between tRF-1 and tRNA^Leu^(TTA). In other words, tRF-1 likely promotes cell death through complementary pathways to the downregulation of mature tRNA^Leu^(TAA).

Based on our results, we propose a model for cell death induced by DDAs that occurs in an SLFN11-dependent manner (Figure 5). CPT, a DDA, induces the downregulation of tRNA^Leu^(TAA) and the production of tRNA fragments (tRFs) in an SLFN11-dependent manner. In this process, the downregulation of tRNA^Leu^(TAA) induces ER stress, leading to activation of the IRE1α‒JNK pathway, CHOP expression, and subsequent cell death. In this case, translation errors at the UUA codon may lead to abnormal proteolysis, the production of unfolded proteins, and mitochondrial dysfunction, resulting in ER stress and disruption of proteostasis. In parallel, tRFs induce cell death by activating the IRE1α‒JNK pathway, thereby promoting CPT-induced cell death.

**Figure 5:**
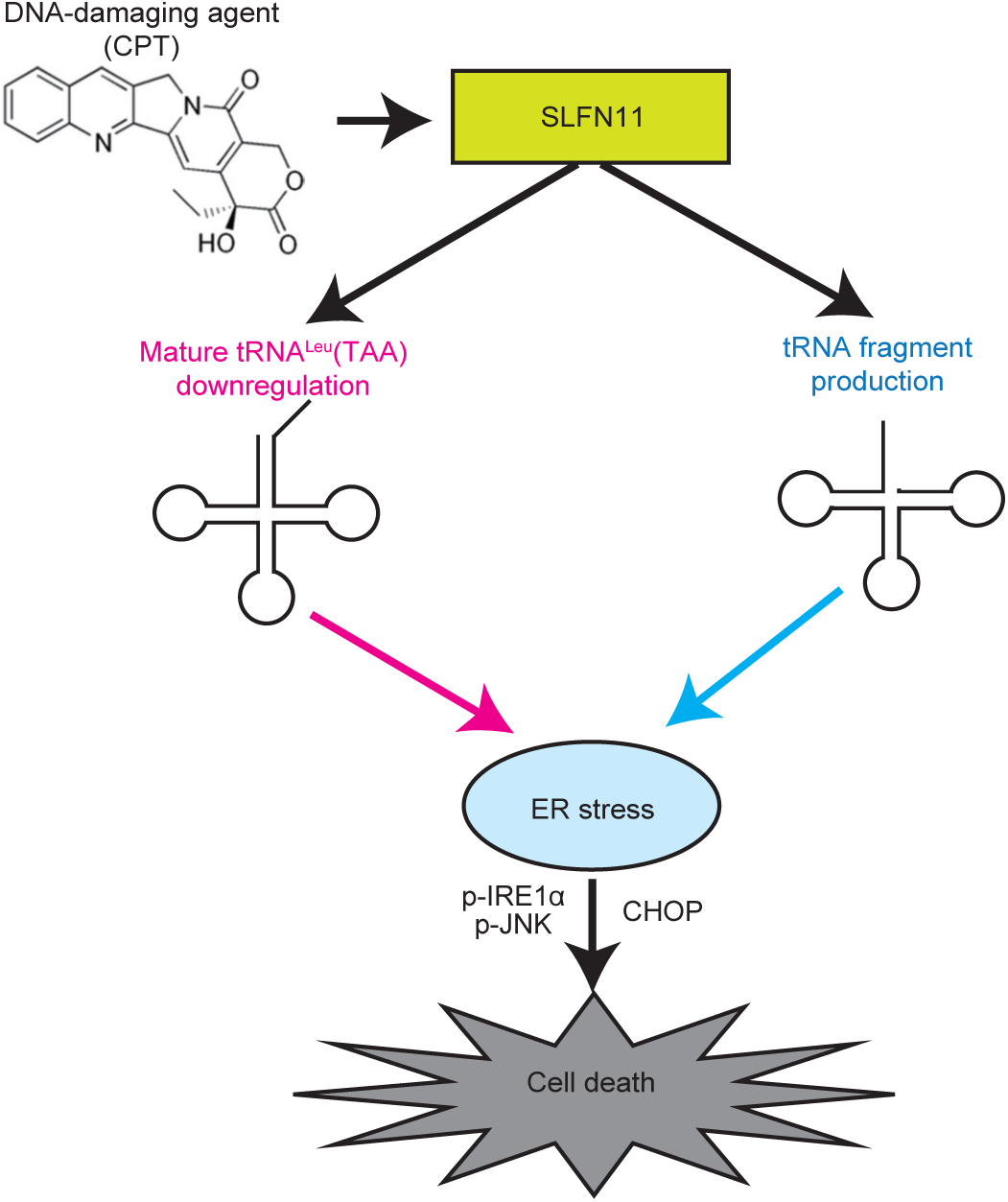
Schematic of CPT-induced cell death via SLFN11-dependent downregulation of tRNA^Leu^(TAA) and tRF production. CPT, a DNA-damaging agent (DDA), induces the downregulation of tRNA^Leu^(TAA) and the production of tRNA fragments in an SLFN11-dependent manner. In this process, downregulation of tRNA^Leu^(TAA) induces ER stress, leading to activation of the IRE1α‒JNK pathway, induction of CHOP expression, and subsequent cell death. In this case, translation errors at the UUA codon may lead to abnormal proteolysis, the production of unfolded proteins, and mitochondrial dysfunction, resulting in ER stress and disruption of proteostasis. In parallel, tRNA fragments (tRFs) induce cell death by activating the IRE1α‒JNK pathway and promote CPT-induced cell death.

## DISCUSSION

SLFN11 sensitizes cancer cells to DDAs. In this study, we demonstrated that SLFN11-dependent downregulation of tRNA^Leu^(TTA) induces ER stress and disrupts proteostasis during CPT treatment, ultimately inducing cell death. Additionally, we showed that specific tRNA fragments (tRF-1 and −2) generated in this process promote CPT-induced cell death.

To date, regarding the involvement of the SLFN family in proteostasis, *SLFN11*-KO cells are more sensitive to a ubiquitin-activating enzyme inhibitor than the parent cells, suggesting SLFN11 has potential to attenuate ER stress (37). In comparison, our findings indicate that SLFN11 induces ER stress under DDA treatment (Figure 2). This difference may arise because, while ubiquitin-activating enzyme inhibitors directly induce protein toxicity, DDAs induce protein toxicity by downregulating tRNA^Leu^(TAA) in a SLFN11-dependent manner. Additionally, a missense mutation in *SLFN2* has been shown to cause chronic ER stress (38). SLFN2 binds to tRNAs but lacks endoribonuclease activity, protecting tRNAs from stress-induced cleavage (39). SLFN2 may play a role in maintaining tRNA^Leu^(TAA) levels and attenuating ER stress.

Regarding the cause of ER stress and disruption of proteostasis, bioinformatics analysis indicated that the reduction of tRNA^Leu^(TAA) may affect the translation of proteins involved in proteolysis, protein transport, and the mitochondria respiratory chain due to the high UUA codon usage frequency in these proteins (Figure 3). Translation errors at UUA codons could lead to the production of unfolded or aggregated proteins (40–42). Furthermore, abnormalities in proteolysis, protein transport, and mitochondrial functions could cause the additional accumulation of unfolded proteins through impaired proteolytic activity or increased production of reactive oxygen species (19). Although further investigation is needed to validate this hypothesis, it is supported by previous studies showing that SLFN12 downregulates tRNA^Leu^(TAA) and causes ribosomal stalling at the UUA codon on mRNAs associated with proteostasis, including ubiquitination and mitochondrial respiratory chain complex I (16).

Although SLFN11-dependent cell death was reported to result from either the ablation of ATR protein with high UUA codon usage frequency (11), or ribotoxic stress signaling via ZAKα during ribosome stalling at UUA codons (17), the cell death observed in our study was induced via ER stress and regulated by the UPR sensor IRE1α (Figure 2). The pathways leading to cell death following tRNA^Leu^(TAA) downregulation may vary depending on cell type and stress intensity (e.g., drug concentration), because the cellular stress response involves various feedback mechanisms to maintain homeostasis (21,43). Previous studies have shown that UPR activation is associated with resistance to DDAs in solid cancers and that IRE1α inhibitors enhance sensitivity to DDAs (44,45). Therefore, ER stress induced by SLFN11-mediated tRNA regulation may be a key factor driving cell death in solid cancers.

This study also provides new insight into how specific tRFs (tRF-1 and −2) produced by SLFN11 induce cell death (Figure 4). In particular, the activation of the IRE1α‒JNK pathway by tRF-1 (Figure 4H) supports the idea that tRF-1 is directly involved in CPT-induced cell death. However, tRFs were also present in the condition where full-length tRNA^Leu^(TAA) rescued cancer cells from apoptosis (Figure 1), suggesting that these regulatory mechanisms may be complex. Furthermore, it is possible that the *in vitro*-synthesized tRFs used in this study were recognized as foreign RNA, inducing cell death via an immune response (46,47). Therefore, further investigations are needed to clarify the relationship between tRF-1 and −2 and the induction of apoptosis observed in this study.

In conclusion, this study demonstrates that intracellular proteostasis regulated by tRNA^Leu^(TAA) plays an important role in cancer cell death caused by DNA-damaging agents. This finding is expected to enhance our understanding of the mechanism underlying drug resistance to DDAs and further contribute to the development of novel cancer therapies targeting SLFN11-mediated tRNA regulation.

## DATA AVAILABILITY

Raw proteomic data files are deposited in jPOST (48) (http://jpostdb.org; jPOST ID: JPST003122/PXD052458, https://repository.jpostdb.org/preview/2040952864664d6cca44bb5) with accession code 7792. The data underlying this article will be shared on reasonable request to the corresponding author.

## SUPPLEMENTARY DATA

Supplementary Data are available at online.

## AUTHOR CONTRIBUTIONS

Conceptualization: Y.I., and A.K.; Methodology: Y.I., T.Mo., T.Ma., S.Ki., N.K., S.Ka., J.G., A.T.S, and A.K.; Investigation and data analysis: Y.I., T.Mo., T.Ma., S.Ki., N.K., S.Ka. and J.G.; Writing – original draft: Y.I. and A.K.; Writing – review & editing: Y.I., T.Mo., T.Ma., S.Ki., N.K., S.Ka., J.G., A.T.S, and A.K.

## ACKNOWLEDGEMENTS

We thank Junko Murai for providing us with the *SLFN11*-KO TOV-112D cells. We thank Kohei Fujiwara for discussions about this study. We also thank Lan Anh Catherine Nguyen and Shunsuke Seo for providing technical support. The graphical abstract was Created in BioRender. Iimori, Y. (2025) https://BioRender.com/x65o789.

## FUNDING

This work was supported by the Japan Society for the Promotion of Science KAKENHI [JP22K16773 and JP24K19705 to Y.I., 23K04996 to T.Mo., 24K10318 to S.Ki.]; the Research Support Project for Life Science and Drug Discovery (Basis for Supporting Innovative Drug Discovery and Life Science Research [BINDS]) from Japan Agency for Medical Research and Development [23ama121018 to T.Ma.]; the Japan Agency for Medical Research and Development program [22ama221112h0001 to A.T.S.]; the National Institutes of Health grants [R01GM144426 to A.T.S.]; and the research funds from the Yamagata prefectural government and the City of Tsuruoka [to Y.I., T.Mo., T.Ma., S.Ki., N.K., S.Ka., J.G., A.T.S., and A.K.]. Funding for open access charge: the Japan Society for the Promotion of Science KAKENHI/JP24K19705.

## CONFLICT OF INTEREST

None declared.

## Notes

### Competing Interest Statement

The authors have declared no competing interest.

