## Supplementary Data for "SLFN11-mediated tRNA regulation induces cell death by disrupting proteostasis in response to DNA-damaging agents"

Supplementary Figures S1 – S6

Supplementary Tables S1 – S6

### MANUSCRIPT TITLE

SLFN11-mediated tRNA regulation induces cell death by disrupting proteostasis in response to DNA-damaging agents

### AUTHORS

Yuki Iimori[1,2], Teppei Morita[1,2], Takeshi Masuda[1,2], Shojiro Kitajima[1,2], Nobuaki Kono[1,2], Shun Kageyama[1,2], Josephine Galipon[1,2,3], Atsuo T. Sasaki[1,2,4,5,6,7], and Akio Kanai[1,2]*

[1] Institute for Advanced Biosciences, Keio University, Tsuruoka, 997-0017, Japan

[2] Systems Biology Program Graduate School of Media and Governance, Keio University, Fujisawa, 252-8520, Japan

[3] Graduate School of Science and Engineering, Yamagata University, Yonezawa, 992-8510, Japan

[4] Division of Hematology and Oncology, Department of Internal Medicine, University of Cincinnati College of Medicine, Cincinnati, OH, 45267, USA

[5] Department of Cancer Biology, University of Cincinnati College of Medicine, Cincinnati, OH, 45267, USA

[6] Department of Neurosurgery, Brain Tumor Center at UC Gardner Neuroscience Institute, Cincinnati, Cincinnati, OH, 45267, USA

[7] Department of Clinical and Molecular Genetics, Hiroshima University Hospital, Hiroshima, 734-8551, Japan.

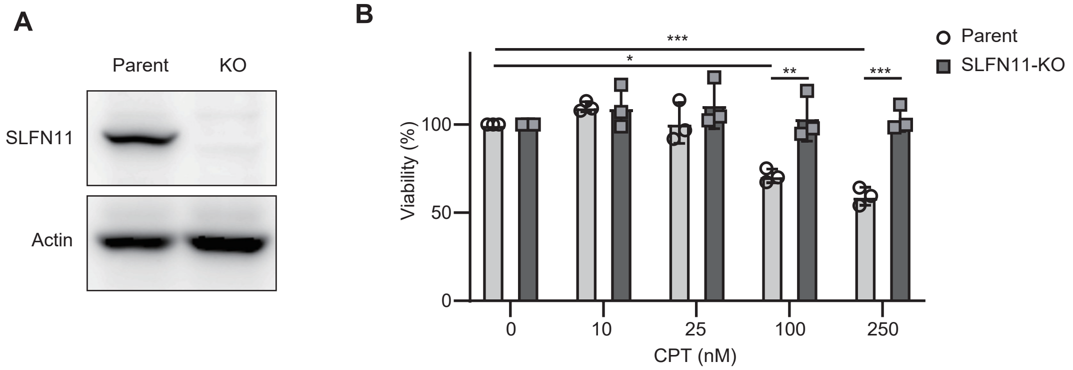


**Supplementary Figure S1 | CPT induces SLFN11-dependent cell death in human cancer cells.** (**A**) Western blotting analysis of SLFN11 protein expression in TOV-112D parent cells and SLFN11-KO cells. Actin was used as a loading control. (**B**) Cell viability of TOV-112D parent cells and SLFN11-KO cells treated with the indicated concentrations of CPT for 24 hours. Data are shown as the mean ± SD (three biological replicates). Two-way ANOVA with Šídák's multiple comparison test was used. *$P$ < 0.05, **$P$ < 0.01, and ***$P$ < 0.001.

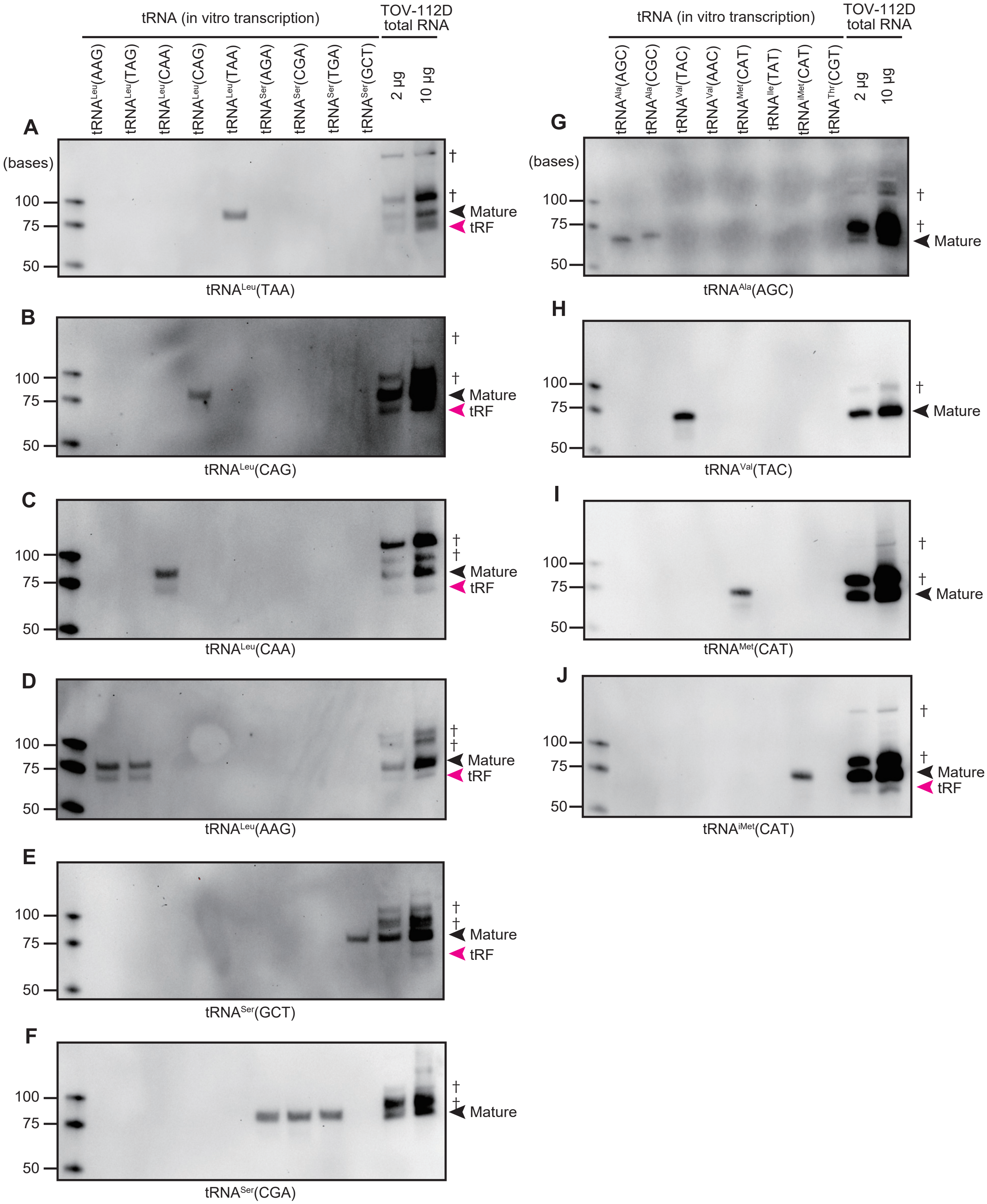
tRNA (in vitro transcription)
TOV-112D total RNA
tRNALeu(AAG)
tRNALeu(TAG)
tRNALeu(CAA)
tRNALeu(CAG)
tRNALeu(TAA)
tRNASer(AGA)
tRNASer(CGA)
tRNASer(TGA)
tRNASer(GCT)
2 µg
10 µg
A
(bases)
100
75
50
†
Mature
tRF
tRNALeu(TAA)
B
tRNALeu(CAG)
C
tRNALeu(CAA)
D
tRNALeu(AAG)
E
tRNASer(GCT)
F
tRNASer(CGA)
tRNAAla(AGC)
tRNAAla(CGC)
tRNAVal(TAC)
tRNAVal(AAC)
tRNAMet(CAT)
tRNAIle(TAT)
tRNAiMet(CAT)
tRNAThr(CGT)
G
tRNAAla(AGC)
H
tRNAVal(TAC)
I
tRNAMet(CAT)
J
tRNAiMet(CAT)

**Supplementary Figure S2 | Northern blotting assessment of RNA probe specificity.** (**A-J**) Northern blotting assessment of the specificities of the RNA probes for each tRNA. The target tRNAs are indicated below each figure. The samples comprised tRNAs synthesized *in vitro* and total RNAs obtained from TOV-112D cells (2 or 10 μg) after 10 hours of treatment with CPT (100 nM). The cross-reactivities of the RNA probes for type II tRNAs ($tRNA^{Leu}$ or $tRNA^{Ser}$) were evaluated. The cross-reactivities of the RNA probes for type I tRNAs including $tRNA^{Ala}$(AGC), $tRNA^{Val}$(TAC), $tRNA^{Met}$(CAT) and $tRNA^{iMet}$(CAT) were evaluated against the tRNA with the highest probability of cross-reacting according to the free energy calculated by IntaRNA (Ref. 1-4) based on the detection region. The black arrowhead, red arrowhead, and dagger indicate the location of mature tRNA, tRF, and the presumed precursor tRNA, respectively.

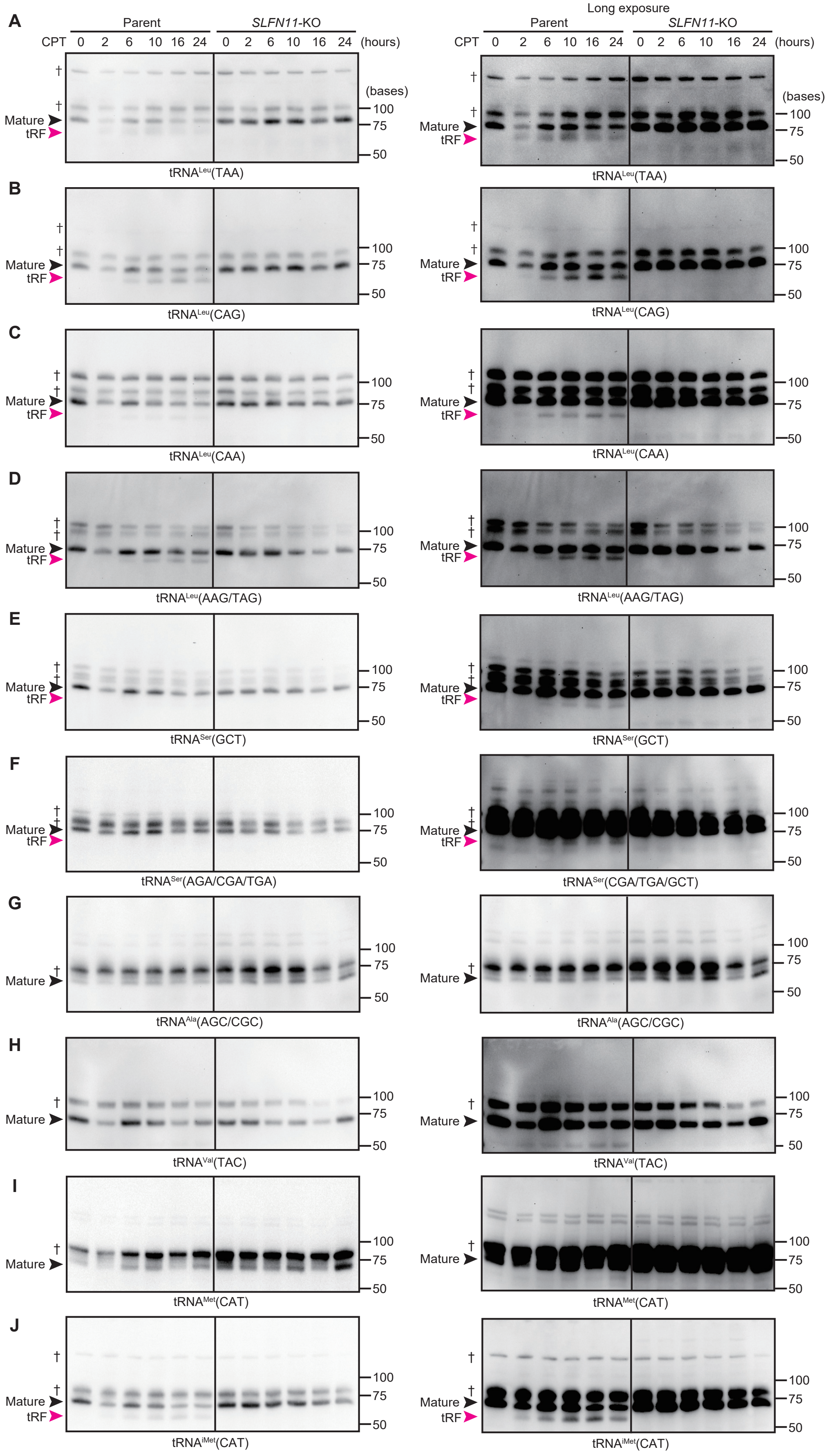
A
Long exposure
Parent
SLFN11-KO
CPT 0 2 6 10 16 24 0 2 6 10 16 24 (hours)
(bases)
100
75
50
†
Mature
tRF
tRNALeu(TAA)
B
tRNALeu(CAG)
C
tRNALeu(CAA)
D
tRNALeu(AAG/TAG)
E
tRNASer(GCT)
F
tRNASer(AGA/CGA/TGA)
tRNASer(CGA/TGA/GCT)
G
tRNAAla(AGC/CGC)
H
tRNAVal(TAC)
I
tRNAMet(CAT)
J
tRNAiMet(CAT)

**Supplementary Figure S3 | Time-series analysis of tRNA level and tRNA fragmentation during CPT treatment.** (**A-J**) Northern blotting of the expression levels of the indicated mature tRNAs (black arrowhead) and tRFs (red arrowhead). TOV-112D parent cells and SLFN11-KO cells were treated with CPT (100 nM) for the indicated time. The dagger indicates the location of the presumed precursor tRNA. Long exposure data from the left column are shown in the right column. The marker size is indicated on the right side. Representative results from two independent experiments are shown.

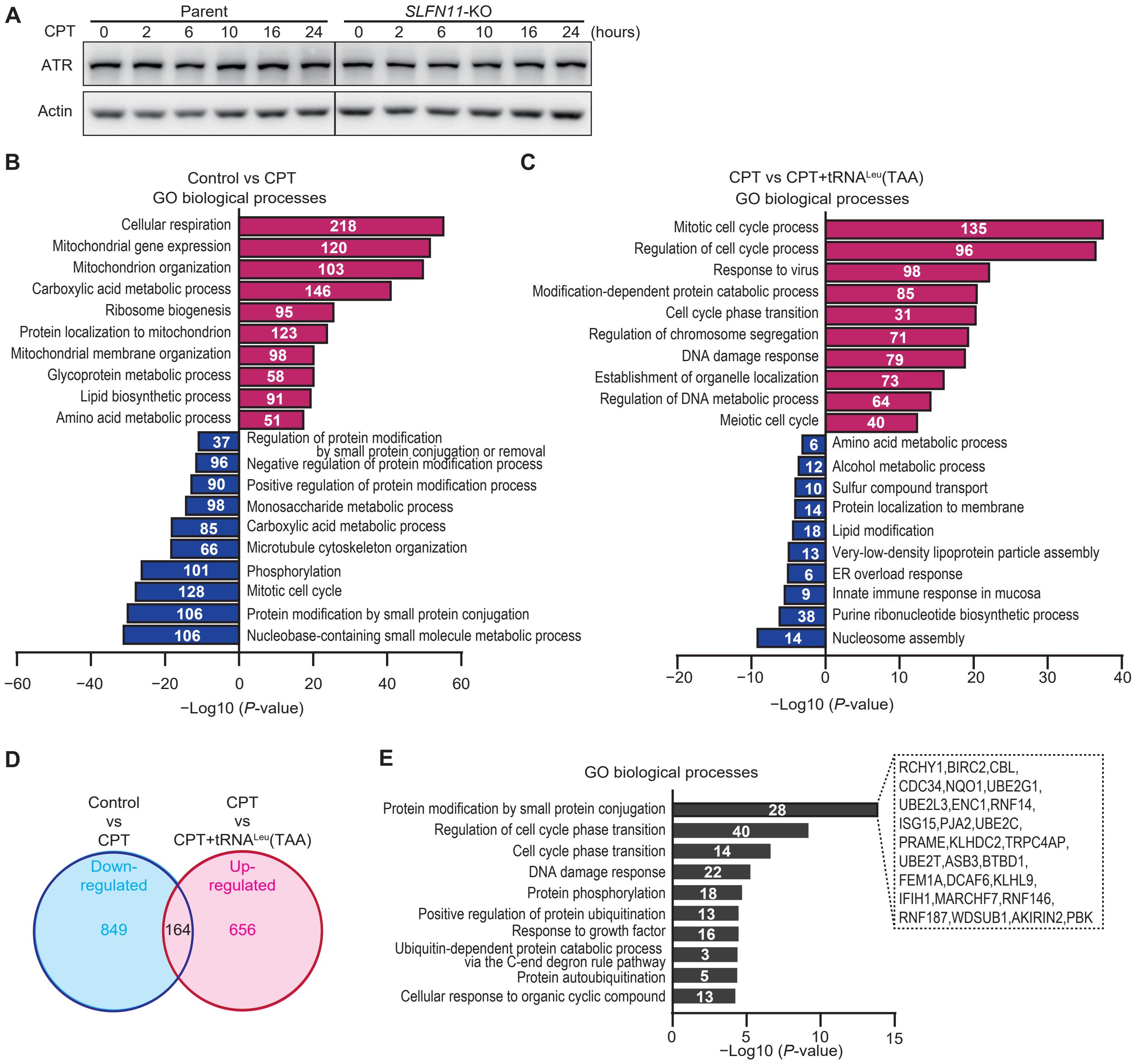

A
CPT
Parent
SLFN11-KO
0 2 6 10 16 24 0 2 6 10 16 24 (hours)
ATR
Actin
B
Control vs CPT
GO biological processes
Cellular respiration 218
Mitochondrial gene expression 120
Mitochondrion organization 103
Carboxylic acid metabolic process 146
Ribosome biogenesis 95
Protein localization to mitochondrion 123
Mitochondrial membrane organization 98
Glycoprotein metabolic process 58
Lipid biosynthetic process 91
Amino acid metabolic process 51
37 Regulation of protein modification by small protein conjugation or removal
96 Negative regulation of protein modification process
90 Positive regulation of protein modification process
98 Monosaccharide metabolic process
85 Carboxylic acid metabolic process
66 Microtubule cytoskeleton organization
101 Phosphorylation
128 Mitotic cell cycle
106 Protein modification by small protein conjugation
106 Nucleobase-containing small molecule metabolic process
−60 −40 −20 0 20 40 60
−Log10 (P-value)
C
CPT vs CPT+tRNALeu(TAA)
GO biological processes
Mitotic cell cycle process 135
Regulation of cell cycle process 96
Response to virus 98
Modification-dependent protein catabolic process 85
Cell cycle phase transition 31
Regulation of chromosome segregation 71
DNA damage response 79
Establishment of organelle localization 73
Regulation of DNA metabolic process 64
Meiotic cell cycle 40
6 Amino acid metabolic process
12 Alcohol metabolic process
10 Sulfur compound transport
14 Protein localization to membrane
18 Lipid modification
13 Very-low-density lipoprotein particle assembly
6 ER overload response
9 Innate immune response in mucosa
38 Purine ribonucleotide biosynthetic process
14 Nucleosome assembly
−20 −10 0 10 20 30 40
−Log10 (P-value)
D
Control vs CPT
CPT vs CPT+tRNALeu(TAA)
Down-regulated
Up-regulated
849
164
656
E
GO biological processes
Protein modification by small protein conjugation 28
Regulation of cell cycle phase transition 40
Cell cycle phase transition 14
DNA damage response 22
Protein phosphorylation 18
Positive regulation of protein ubiquitination 13
Response to growth factor 16
Ubiquitin-dependent protein catabolic process via the C-end degron rule pathway 3
Protein autoubiquitination 5
Cellular response to organic cyclic compound 13
0 5 10 15
−Log10 (P-value)
RCHY1,BIRC2,CBL,
CDC34,NQO1,UBE2G1,
UBE2L3,ENC1,RNF14,
ISG15,PJA2,UBE2C,
PRAME,KLHDC2,TRPC4AP,
UBE2T,ASB3,BTBD1,
FEM1A,DCAF6,KLHL9,
IFIH1,MARCHF7,RNF146,
RNF187,WDSUB1,AKIRIN2,PBK

**Supplementary Figure S4 | Downregulation of tRNA$^{Leu}$(TAA) induces ER stress.** (**A**) Western blotting analysis of the time-series of ATR protein expression after treating TOV-112D parent cells and SLFN11-KO cells with CPT (100 nM). Actin was used as a loading control. Representative results from two independent experiments are shown. (**B** and **C**) Enrichment analysis of Figure 2A and 2B are shown in (B) and (C), respectively. The results for significantly upregulated proteins are shown on the right (red bars) and significantly downregulated proteins are shown on the left (blue bars). The white numbers in the bars indicate the number of proteins in each group. (**D**) Venn diagram of the overlap between downregulated proteins in Figure 2A and upregulated proteins in Figure 2B. The total number of overlapping proteins is shown in the center in black letters. (**E**) Enrichment analysis of the overlapping proteins in (D). The white numbers in the bars indicate the number of proteins in each group. The proteins involved in the protein modification by small protein conjugation are annotated on the right of the bar.

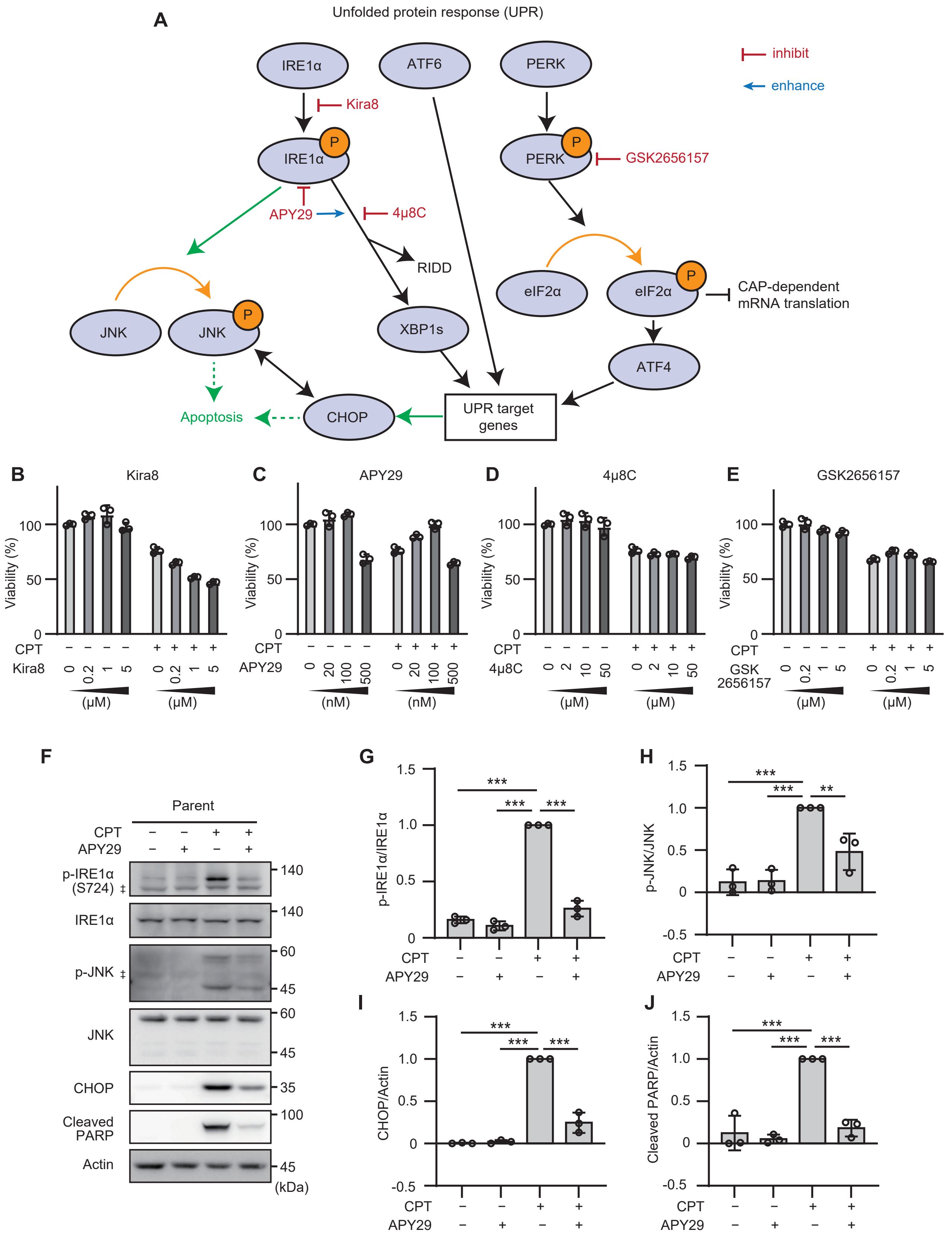
A
Unfolded protein response (UPR)
IRE1α
ATF6
PERK
inhibit
enhance
Kira8
P
GSK2656157
APY29
4μ8C
RIDD
eIF2α
CAP-dependent
mRNA translation
JNK
XBP1s
ATF4
Apoptosis
CHOP
UPR target
genes
B
Kira8
C
APY29
D
4μ8C
E
GSK2656157
Viability (%)
CPT
GSK
2656157
(μM)
(nM)
F
Parent
CPT
APY29
p-IRE1α
(S724)
IRE1α
p-JNK
JNK
CHOP
Cleaved
PARP
Actin
(kDa)
G
p-IRE1α/IRE1α
H
p-JNK/JNK
I
CHOP/Actin
J
Cleaved PARP/Actin

**Supplementary Figure S5 | CPT induces cell death via ER stress.** (**A**) Schematic of the Unfolded protein response（UPR) pathways and the inhibitory sites of each inhibitor (Ref. 5 & 6). (**B-E**) Cell viability of TOV-112D cells was assessed after 24 hours of treatment with CPT (100 nM) and the indicated concentrations of the stress senser inhibitors: Kira8 (B), APY29 (C), 4μ8c (D), and GSK265615 (E). Values are relative to the mean value of control samples treated with 0.1% DMSO. Data are shown as the mean ± SD (three technical replicates). (**F**) Western blotting analysis of the UPR after treatment of TOV-112D cells with CPT (100 nM) and/or APY29 (100 nM) for 24 hours. Representative results of three independent experiments are shown. Double daggers indicate the position of the non-specific bands. The marker size is indicated on the right side. (**G-J**) Quantification of the protein expression levels of phospho-IRE1α (G), phospho-JNK (H), CHOP (I) and cleaved PARP (J) shown in (F). Data were normalized to the expression level of IRE1α (G), JNK (H), and actin (I and J) in the same sample. Values are relative to the values of the samples treated with CPT (100 nM). Data are shown as the mean ± SD (three biological replicates). One-way ANOVA with Dunnett's multiple comparison test was used. **P* < 0.05, ***P* < 0.01, and ****P* < 0.001.

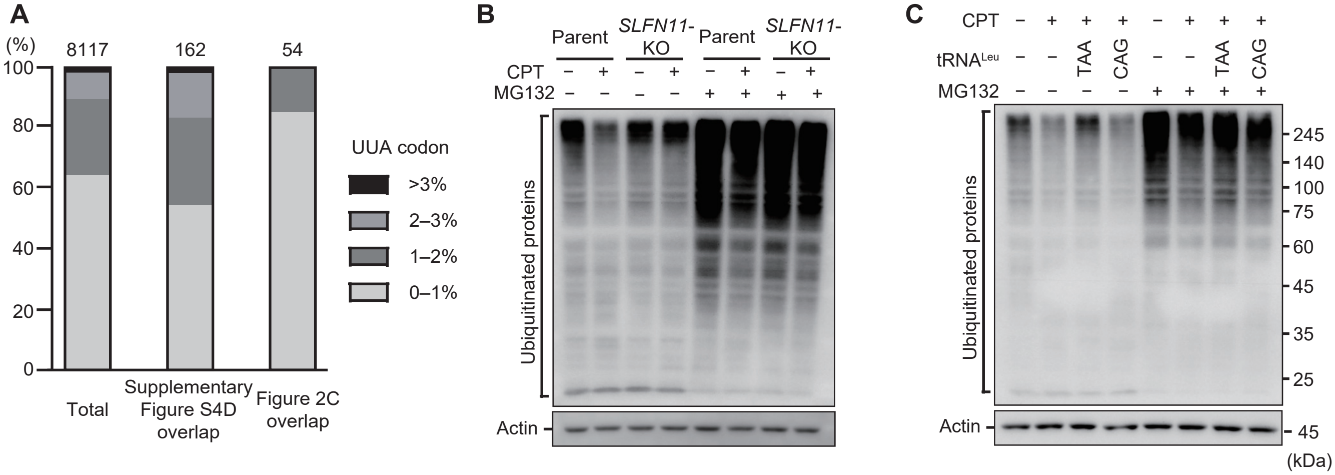


**Supplementary Figure S6 | tRNA$^{Leu}$(TAA) has the potential to regulate proteostasis through translation.** (**A**) Proportion of proteins with UUA codon usage frequency in the indicated range. The left bar shows all detected proteins (8117 proteins), the middle bar shows the overlapping proteins from Supplementary Figure S4D (162 proteins), and the right bar shows the overlapping proteins from Figure 2C (54 proteins). (**B** and **C**) Western blotting analysis of the ubiquitinated proteins in TOV-112D parent cells or *SLFN11*-KO cells treated with CPT (100 nM) for 24 hours (B) or parent cells transfected with the indicated tRNA (10 nM) followed by treatment with CPT (100 nM) for 24 hours (C). The samples in the right-hand lanes were treated with MG132 (5 μM) from 18 to 24 hours after CPT administration. Representative results from three independent experiments are shown. Actin was used as a loading control. The marker size is indicated on the right side.

**Supplementary Table S1. Antisense RNA probes.**

| RNA probe name | Sequence | Target portion of RNA probe* (reference gene†) |
|---|---|---|
| **tRNA$^{\text{Leu}}$(TAA)-probe** | 5′-GGUUCGAACCCACGCGGACAUAUGUCCAUUGGAUCUUAAG-3′ | Nucleotides 32–61 of tRNA$^{\text{Leu}}$(TAA) (tRNA-Leu-TAA-1-1) |
| **tRNA$^{\text{Leu}}$(CAG)-probe** | 5′-GCACGCCUCCAGGGGAGACUGCGACCUGAACGCAGC-3′ | Nucleotides 26–51 of tRNA$^{\text{Leu}}$(CAG) (tRNA-Leu-CAG-1-1) |
| **tRNA$^{\text{Leu}}$(CAA)-probe** | 5′-GCACGCCUCCAUUGGAGACCAGAACUUGAGUCUGGC-3′ | Nucleotides 26–51 of tRNA$^{\text{Leu}}$(CAA) (tRNA-Leu-CAA-1-1) |
| **tRNA$^{\text{Leu}}$(AAG)-probe** | 5′-GCACGCCUCCGAAGAGACUGGAGCCUUAAUCCAGCG-3′ | Nucleotides 25–51 of tRNA$^{\text{Leu}}$(AAG) (tRNA-Leu-AAG-1-1) |
| **tRNA$^{\text{Ser}}$(GCT)-probe** | 5′-GAUUCGAACCCACGCGUGCAGAGCACAAUGGAUUAGCAGU-3′ | Nucleotides 31–61 of tRNA$^{\text{Ser}}$(GCT) (tRNA-Ser-GCT-1-1) |
| **tRNA$^{\text{Ser}}$(CGA)-probe** | 5′-GCGGGGAGACCCCAUUGGAUUUCGAGUCCAACGCC-3′ | Nucleotides 22–48 of tRNA$^{\text{Ser}}$(CGA) (tRNA-Ser-CGA-1-1) |
| **tRNA$^{\text{Ala}}$(AGC)-probe** | 5′-GCCAGGACCUCGUGCAUGCUAAGCACGCGCUCUACC-3′ | Nucleotides 18–52 of tRNA$^{\text{Ala}}$(AGC) (tRNA-Ala-AGC-1-1) |
| **tRNA$^{\text{Val}}$(TAC)-probe** | 5′-GCCAGGACCUUCUGCGUGUAAAGCAGACGUGAUAAC-3′ | Nucleotides 19–52 of tRNA$^{\text{Val}}$(TAC) (tRNA-Val-TAC-1-1) |
| **tRNA$^{\text{Met}}$(CAT)-probe** | 5′-GACUCACGACCUUCAGAUUAUGAGACUGACGCGCUA-3′ | Nucleotides 20–54 of tRNA$^{\text{Met}}$(CAT) (tRNA-Met-CAT-1-1) |
| **tRNA$^{\text{iMet}}$(CAT)-probe** | 5′-GACCUCUGGGUUAUGGGCCCAGCACGCUUCCGCU-3′ | Nucleotides 14–48 of tRNA$^{\text{iMet}}$(CAT) (tRNA-iMet-CAT-1-1) |

The "G" nucleotide that was artificially added for efficient transcription from the T7 promoter is underlined. *The target portions of the RNA probes are summarized according to the universal tRNA positioning rules (Ref. 7). †Reference genes are shown according to GtRNAdb gene symbol (Refs. 8 & 9).

**Supplementary Table S2. tRNA transcripts.**

| tRNA name | Sequence | Location of the transcribed tRNA* (reference gene†) |
|---|---|---|
| **Full-length tRNA** | | Nucleotides 1–76 of: 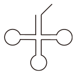 |
| **Full-length tRNA$^{Leu}$(TAA)** | 5′-GACCAGGAUGGCCGAGUGGUUAAGGCGU UGGACUUAAGAUCCAAUGGACAUAUGUCCG CGUGGGUUCGAACCCCACUCCUGGUACCA-3′ | tRNA$^{Leu}$(TAA) (tRNA-Leu-TAA-1-1) |
| **Full-length tRNA$^{Leu}$(AAG)** | 5′-GGUAGCGUGGCCGAGCGGUcUAAGGCGC UGGAUUAAGGCUCCAGUCUCUUCGGAGGCG UGGGUUCGAAUCCCACCGCUGCCACCA-3′ | tRNA$^{Leu}$(AAG) (tRNA-Leu-AAG-1-1) |
| **Full-length tRNA$^{Leu}$(TAG)** | 5′-GGUAGCGUGGCCGAGCGGUcUAAGGCGC UGGAUUUAGGCUCCAGUCUCUUCGGAGGCG UGGGUUCGAAUCCCACCGCUGCCACCA-3′ | tRNA$^{Leu}$(TAG) (tRNA-Leu-TAG-1-1) |
| **Full-length tRNA$^{Leu}$(CAA)** | 5′-GUCAGGAUGGCCGAGUGGUcUAAGGCGC CAGACUCAAGUUCUGGUCUCCAAUGGAGGC GUGGGUUCGAAUCCCACUUCUGACACCA-3′ | tRNA$^{Leu}$(CAA) (tRNA-Leu-CAA-1-1) |
| **Full-length tRNA$^{Leu}$(CAG)** | 5′-GUCAGGAUGGCCGAGCGGUcUAAGGCGC UGCGUUCAGGUCGCAGUCUCCCCUGGAGGC GUGGGUUCGAAUCCCACUCCUGACACCA-3′ | tRNA$^{Leu}$(CAG) (tRNA-Leu-CAG-1-1) |
| **Full-length tRNA$^{Ser}$(AGA)** | 5′-GUAGUCGUGGCCGAGUGGUUAAGGCGAU GGACUAGAAAUCCAUUGGGGUUUCCCCGCG CAGGUUCGAAUCCUGCCGACUACGCCA-3′ | tRNA$^{Ser}$(AGA) (tRNA-Ser-AGA-1-1) |
| **Full-length tRNA$^{Ser}$(CGA)** | 5′-GCUGUGAUGGCCGAGUGGUUAAGGCGUU GGACUCGAAAUCCAAUGGGGUCUCCCCGCG CAGGUUCGAAUCCUGCUCACAGCGCCA-3′ | tRNA$^{Ser}$(CGA) (tRNA-Ser-CGA-1-1) |
| **Full-length tRNA$^{Ser}$(TGA)** | 5′-GCAGCGAUGGCCGAGUGGUUAAGGCGUU GGACUUGAAAUCCAAUGGGGUCUCCCCGCG CAGGUUCGAACCCUGCUCGCUGCGCCA-3′ | tRNA$^{Ser}$(TGA) (tRNA-Ser-TGA-1-1) |
| **Full-length tRNA$^{Ser}$(GCT)** | 5′-GACGAGGUGGCCGAGUGGUUAAGGCGAU GGACUGCUAAUCCAUUGUGCUCUGCACGCG UGGGUUCGAAUCCCACCCUCGUCGCCA-3′ | tRNA$^{Ser}$(GCT) (tRNA-Ser-GCT-1-1) |
| **Full-length tRNA$^{Ala}$(AGC)** | 5′-GGGGGUAUAGCUCAGUGGUAGAGCGCGU GCUUAGCAUGCACGAGGUCCUGGGUUCGAU CCCCAGUACCUCCACCA-3′ | tRNA$^{Ala}$(AGC) (tRNA-Ala-AGC-1-1) |
| **Full-length tRNA$^{Ala}$(CGC)** | 5′-GGGGGUGUAGCUCAGUGGUAGAGCGCGU GCUUCGCAUGUACGAGGcCCCGGGUUCGAC CCCCGGCUCCUCCACCA-3′ | tRNA$^{Ala}$(CGC) (tRNA-Ala-CGC-4-1) |
| **Full-length tRNA$^{Val}$(TAC)** | 5′-GGUUCCAUAGUGUAGUGGUUAUCACGUC UGCUUUACACGCAGAAGGUCCUGGGUUCGA GCCCCAGUGGAACCACCA-3′ | tRNA$^{Val}$(TAC) (tRNA-Val-TAC-1-1) |

| | | | |
|---|---|---|---|
| **Full-length tRNA$^{Val}$(AAC)** | 5′-GUUUCCGUAGUGUAGUGGUUAUCACGUU UGCCUAACACGCGAAAGGUCCCCGGUUCGA AACCGGGCAGAAACACCA-3′ | tRNA$^{Val}$(AAC) (tRNA-Val-AAC-5-1) | |
| **Full-length tRNA$^{Met}$(CAT)** | 5′-GCCUCGUUAGCGCAGUAGGUAGCGCGUC AGUCUCAUAAUCUGAAGGUCGUGAGUUCGA UCCUCACACGGGGCACCA-3′ | tRNA$^{Met}$(CAT) (tRNA-Met-CAT-1-1) | |
| **Full-length tRNA$^{Ile}$(TAT)** | 5′-GCUCCAGUGGCGCAAUCGGUUAGCGCGC GGUACUUAUAAUGCCGAGGUUGUGAGUUCG AGCCUCACCUGGAGCACCA-3′ | tRNA$^{Ile}$(TAT) (tRNA-Ile-TAT-2-1) | |
| **Full-length tRNA$^{iMet}$(CAT)** | 5′-GAGCAGAGUGGCGCAGCGGAAGCGUGCU GGGCCCAUAACCCAGAGGUCGAUGGAUCGA AACCAUCCUCUGCUACCA-3′ | tRNA$^{iMet}$(CAT) (tRNA-iMet-CAT-1-1) | |
| **Full-length tRNA$^{Thr}$(CGT)** | 5′-GGCCCUGUAGCUCAGCGGUUGGAGCGCU GGUCUCGUAAACCUAGGGGUCGUGAGUUCA AAUCUCACCAGGGCCUCCA-3′ | tRNA$^{Thr}$(CGT) (tRNA-Thr-CGT-5-1) | |
| **tRF-1$^{Leu}$(TAA)** | 5′-GACCAGGAUGGCCGAGUGGUUAAGGCGU UGGACUUAAGAUCCAAUGGACAUAUGUCCG CGUGGGUUCGAACCCCACU-3′ | Nucleotides 1–66 of tRNA$^{Leu}$(TAA) (tRNA-Leu-TAA-1-1) | 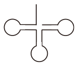 |
| **tRF-2$^{Leu}$(TAA)** | 5′-GCUGCACCAGGAUGGCCGAGUGGUUAAG GCGUUGGACUUAAGAUCCAAUGGACAUAUG UCCGCGU-3′ | Nucleotides −4–50 of tRNA$^{Leu}$(TAA) (tRNA-Leu-TAA-1-1) | 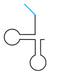 |
| **tRF-3$^{Leu}$(TAA)** | 5′-GAUGGCCGAGUGGUUAAGGCGUUGGACU UAAGAUCCAAUGGACAUAUGUCCGCGUGGG UUCGAAC-3′ | Nucleotides 6–60 of tRNA$^{Leu}$(TAA) (tRNA-Leu-TAA-1-1) | 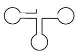 |

The "G" nucleotide that was artificially added for efficient transcription from the T7 promoter is underlined. *The locations of the transcribed tRNAs are summarized according to the universal tRNA positioning rules (Ref. 7). †Reference genes are shown according to GtRNAdb gene symbol (Refs. 8 & 9).

**Supplementary Table S3. Primers used to sequence tRNA fragments.**

| Name | Sequence | Comment |
|---|---|---|
| **RT primer_1** | 5′-GCCTTAACCACTCGGCCAT-3′ | RT primer for tRNA$^{Leu}$(TAA) |
| **PCR primer_2** | 5′-TTCCGCCCGCGAAATGATCCAATGGACATATGTCC-3′ | Sense PCR primer for tRNA$^{Leu}$(TAA) cDNA |
| **PCR primer_3** | 5′-CACGAGAATTGCTGGGCCTTAACCACTCGGCCAT-3′ | Antisense PCR primer for tRNA$^{Leu}$(TAA) cDNA |
| **pMiniT_S** | 5′-ATTTCGCGGGCGGAACCC-3′ | Sense PCR primer for pMiniT2.0 plasmid |
| **pMiniT_A** | 5′-CCAGCAATTCTCGTGAATCATCGCT-3′ | Antisense PCR primer for pMiniT2.0 plasmid |

The complementary sequences of vectors for In-Fusion cloning are underlined.

**Supplementary Table S4. List of antibodies used for western blotting.**

| Antibody name (target protein) | Dilution ratio | Company | Product number | Blocking reagent |
|---|---|---|---|---|
| **Primary antibody** | | | | |
| **Pan-actin** | 1:10000 | Cell Signaling Technology, Danvers, MA, USA | #12748 | 4% BSA |
| **IRE1α** (**p Ser724**) | 1:1000 | Novus Biologicals, Centennial, CO, USA | NB100-2323SS | 0.5% skim milk |
| **IRE1α** | 1:500 | Santa Cruz Biotechnology, Dallas, TX, USA | sc-390960 | 0.5% skim milk |
| **Phospho-SAPK/JNK (Thr183/Tyr185)** | 1:1000 | Cell Signaling Technology | #4668 | 0.5% skim milk |
| **JNK** | 1:200 | Santa Cruz Biotechnology | sc-7345 | 0.5% skim milk |
| **CHOP/GADD153** | 1:1000 | Proteintech, Rosemont, IL, USA | 15204-1-AP | 0.5% skim milk |
| **Cleaved PARP** | 1:1000 | Cell Signaling Technology | #5625 | 0.5% skim milk |
| **SLFN11** | 1:2500 | Santa Cruz Biotechnology | sc-515071 | 4% BSA |
| **ATR** | 1:500 | Santa Cruz Biotechnology | sc-515173 | 0.5% skim milk |
| **Ubiquitin** | 1:1000 | Santa Cruz Biotechnology | sc-8017 | 0.5% skim milk |
| **Secondary antibody** | | | | |
| **Anti-mouse IgG, HRP-linked antibody** | 1:5000 | Cell Signaling Technology | #7076 | – |
| **Anti-rabbit IgG, HRP-linked antibody** | 1:5000 | Cell Signaling Technology | #7074 | – |

BSA, bovine serum albumin; HRP, horseradish peroxidase; IgG, immunoglobulin G

**Supplementary Table S5. Inhibitors used in this study and their respective targets.**

| Inhibitor | Target | Company | Product number |
|---|---|---|---|
| **Camptothecin** (**CPT**) | DNA topoisomerase I | Cayman Chemical, Ann Arbor, MI, USA | 11694 |
| **Kira8** | Inositol-requiring enzyme 1 (IRE1α) (type II kinase inhibitor) [Ref. 6] | TargetMol, Boston, MA, USA | T11762L |
| **APY29** | Inositol-requiring enzyme 1 (IRE1α) (type I kinase inhibitor) [Ref. 6] | TargetMol, Boston, MA, USA | T3654 |
| **4μ8C** | Inositol-requiring enzyme 1 (IRE1α) (RNase inhibitor) [Ref. 6] | TargetMol, Boston, MA, USA | T6363 |
| **GSK2656157** | Protein kinase R-like endoplasmic reticulum kinase (PERK) | Selleck, Houston, TX, USA | S7033 |

**Supplementary Table S6. Cleavage sites of tRF$^{Leu}$(TAA).**

| Count | Sequence | Comment |
|---|---|---|
| **7/66** | 5'-(G)ACCAGGATGGCCGAGTGGTtAAGGCGTTGGACTTAAGATCCAATGGACATATGTCCGCGTGGGTTCGAACCCCACTCCTGGTACCA(G)-3' | Full length tRNA |
| **20/66** | 5'- (G)ACCAGGATGGCCGAGTGGTtAAGGCGTTGGACTTAAGATCCAATGGACATATGTCCGCGTGGGTTCGAACCCCACT(G)-3' | tRF-1 |
| **2/66** | 5'- (GG)ACCAGGATGGCCGAGTGGTtAAGGCGTTGGACTTAAGATCCAATGGACATATGTCCGCGTGGGTTCGAACCCCACT(GG)-3' | |
| **6/66** | 5'- (T)ACCAGGATGGCCGAGTGGTtAAGGCGTTGGACTTAAGATCCAATTGACATATGTCCGCGTGGGTTCGAACCCCACT(T)-3' | |
| **2/66** | 5'- (T)CTGCACCAGGATGGCCGAGTGGTtAAGGCGTTGGACTTAAGATCCAATGGACATATGTCCGCG(T)-3' | tRF-2 |
| **2/66** | 5'- (T)CTGCACCAGGACGGCCGAGTGGTtAAGGCGTTGGACTTAAGATCCAATGGACATATGTCCGCG(T)-3' | |
| **2/66** | 5'- (T)CTGCACCACGATGGCCGAGTGGTtAAGGCGTTGGACTTAAGATCCAATGGACATATGTCCGCG(T)-3' | |
| **2/66** | 5'- CTGCACCAGGATGGCCGAGTGGTtAAGGCGTTGGACTTAAGATCCAATGGACATATGTCCGCGTG-3' | |
| **1/66** | 5'-GTCTGCACCAGGATGGCCGAGTGGTtAAGGCGTTGGACTTAAGATCCAATGGACATATGTCCGCG-3' | |
| **2/66** | 5′-GATGGCCGAGTGGTtAAGGCGTTGGACTTAAGATCCAATGGACATATGTCCACGTGGGTTCGAAC-3′ | tRF-3 |
| **1/66** | 5′-GATGGCCGAGTGGTtAAGGCGTTGGACTTAAGATCCAATGGACATATGTCCACGTGGGTCCGAAC-3′ | |
| **1/66** | 5′- (AG)TGGACCAGGATGGCCGAGTGGTtAAGGCGTTGGACTTAAGATCCAATGGACATATGTCCGCGTGGGTTCGAACCCCACT(AG)-3′ | |
| **3/66** | 5′- (ACA)GCACCAGGATGGCCGAGTGGTtAAGGCGTTGGACTTAAGATCCAATGGACATATGTCC(ACA)-3′ | |
| **1/66** | 5′- (G)ACCAGGATGGCCGAGTGGTtAAGGCGTTGGACTTAAGATCCAATGGACATATGTCCGCGTGGGTTCGAAC(G)-3′ | |
| **4/66** | 5′- (G)ATGGCCGAGTGGTtAAGGCGTTGGACTTAAGATCCAATGGACATATGTCCGCGT(G)-3′ | |
| **1/66** | 5′-ATGGCCGAGTGGTtAAGGCGTTGGACTTAAGATCCAATGGACATATGTCCGC-3′ | |
| **9/66** | others | |

tRF$^{Leu}$(TAA) sequences with more than 95% homologous sequences to tRNA$^{Leu}$(TAA). Sequences with less than 95% homology are included in “others”. The read counts of each tRF$^{Leu}$(TAA) are shown in the left column. The representative examples shown in Figure 4A are annotated in the right column. The underlined letters indicate the position of the primers, red letters indicate unexpected sequences, and blue letters indicate the 5′ leader sequences before processing. Bases in brackets are located at either the 5′ end or at the 3′ end. In Figure 4A, the unexpected sequences are omitted.
